# Evolutionary dynamics of genome size and content during the adaptive radiation of Heliconiini butterflies

**DOI:** 10.1101/2022.08.12.503723

**Authors:** Francesco Cicconardi, Edoardo Milanetti, Erika C. Pinheiro de Castro, Anyi Mazo-Vargas, Steven M. Van Belleghem, Angelo Alberto Ruggieri, Pasi Rastas, Joseph Hanly, Elizabeth Evans, Chris D Jiggins, W Owen McMillan, Riccardo Papa, Daniele Di Marino, Arnaud Martin, Stephen H Montgomery

## Abstract

*Heliconius* butterflies, a speciose genus of Müllerian mimics, represent a classic example of an adaptive radiation that includes a range of derived dietary, life history, physiological and neural traits. However, key lineages within the genus, and across the broader Heliconiini tribe, lack genomic resources, limiting our understanding of how adaptive and neutral processes shaped genome evolution during their radiation. We have generated highly contiguous genome assemblies for nine new Heliconiini, 29 additional reference-assembled genomes, and improve 10 existing assemblies. Altogether, we provide a major new dataset of annotated genomes for a total of 63 species, including 58 species within the Heliconiini tribe. We use this extensive dataset to generate a robust and dated heliconiine phylogeny, describe major patterns of introgression, explore the evolution of genome architecture, and the genomic basis of key innovations in this enigmatic group, including an assessment of the evolution of putative regulatory regions at the *Heliconius* stem. Our work illustrates how the increased resolution provided by such dense genomic sampling improves our power to generate and test gene-phenotype hypotheses, and precisely characterize how genomes evolve.

---

A central goal of evolutionary biology is to understand how biodiversity is generated, maintained, and how interactions between organisms drive the diversity of natural communities. Periods of rapid diversification are often associated with the colonisation of new evolutionary niches or the exploitation of new resources^1^. The evolution of key innovations, such as physiological adaptation to food resources, or new morphological traits, can enable these ecological shifts, and play critical roles in adaptive radiations^2^. From a genetic perspective, one fundamental question in understanding how adaptive radiations emerge, is if a significant amount of change, and sources of variability, originate prior to the acceleration in diversification, and whether this variation facilitates the subsequent adaptive radiation. This would be consistent with phyletic gradualism at a genetic level. Identifying and understanding the genetic basis of such key innovations is now a realistic goal^3,4^, and can provide explicit links between genetic changes, natural selection and speciation, in the context of wider patterns of genomic divergence.

Heliconiini, a Neotropical tribe of Nymphalid butterflies, comprised of ∼80 species and ∼400 subspecies, have become a key system to explore the biology of speciation^5–8^. In particular, the rapid radiation of the genus *Heliconius*, and their diversity of colour patterns, have become a case study in how genomic approaches can improve our understanding of the genetic architecture of adaptive traits, and the accumulation of reproductive isolation with ongoing gene flow^7^. Heliconiini also exhibit key innovations including the tribe-wide restricted use of Passifloraceae as larval hostplants – their antagonist coevolutionary partners that can provide them with cyanogenic glucosides for chemical protection^9^. Within the Heliconiini, species of the genus *Heliconius* are also the only lepidoptera to actively collect and digest pollen as adults, which is associated with major shifts in reproductive lifespan^10^, and behavioural and neural elaboration^11^. As such, the availability of tribe-wide genomic resources would represent a major resource to explore the biology of an enigmatic case study in adaptive diversification.

Here, we provide such a resource by sequencing and assembling new genomic data, and using a combinatorial approach to maximise methodological outcomes, with a unified cross-species annotation to remove possible species-biases previously unrepresented in available data. Combined with already available resources, which we also improve both in terms of assembly contiguity and gene annotation, we generate a new genomic dataset that comprises ∼75% of all the species in the Heliconiini tribe, to our knowledge one of the most comprehensive efforts to sample an insect tribe at high taxonomic density. With this, we produce a comprehensive dated phylogeny for Heliconiini and explore patterns of gene flow across the tribe. We test if a significant and substantial amount of genomic change occurred not only at the stem of *Heliconius*, but also at more basal branches within the Heliconiini tribe, pre-dating the range of innovations seen in *Heliconius*. Finally, we investigate structural and adaptive aspects of genome evolution across the radiation and during key ecological transitions, and explore evidence of accelerated evolution in putative regulatory elements, the first analysis of this kind in non-model insect species. Our analyses provide refined views of genomic diversity across Heliconiini, and provide wide ranging new gene-phenotype hypotheses that will provide the foundation for future functional experiments.

## Results & Discussion

### Improved Resolution of Phylogenetic Relationships and Signatures of Introgression Across the Genome

To generate the species tree, we first compiled a total data set of 4,011,390 base pairs of aligned protein-coding DNA obtained from the single-copy orthologous groups (scOGs). The alignment has over 1.5M parsimony-informative, ∼500k singleton sites, and 1.9k constant sites. A species-level phylogeny was determined with a maximum-likelihood (ML) analysis, and used to estimate divergence dates (Fig. 1c, Supplementary Fig. 20, and Supplementary Table 3). Although the topology is widely consistent with previously inferred phylogenetic relationships^8^, we identify differences within some *Heliconius* clades, where the Silvaniform and Melpomene clades are now paraphyletic, and among other genera of Heliconiini, with *Podothricha telesiphe* and *Dryas iulia* now sister lineages, outgrouped by *Dryadula phaetusa* and *Philaethria dido*. The estimated divergence times show that the subfamily Heliconiinae originated ∼45.3 million years (Mya) (95% CI: 35.9-55.5), with the last common ancestors of *Eueides* and *Heliconius* dating to ∼11.1 Mya (95% CI: 7.3-12.1) and 9.6 Mya (95% CI: 8.8-13.8), respectively. Interestingly, deeper branches of the phylogeny are characterized by high molecular substitution rates (Fig. 1c and Supplementary Table 3), indicating a series of bursts in evolutionary rate at the base of the radiation, supported by a highly sampled posterior distribution across our tree (ESS >> 1000; Supplementary Fig. 21). To account for incomplete lineage sorting (ILS) within the phylogeny, we used a coalescent summary method for species trees reconciliation using gene trees (Fig. 2b). This resulted in an almost identical topology as the ML tree (Fig. 1c, Supplementary Fig. 19), with a single exception of the *H. clysonymus + H. hortense + H. telesiphe* branch, which could be due to high rates of ILS or introgression (coalescent units = 0.08), disrupting the monophyly of the Erato clade^12^. We find little evidence of ILS around more basal nodes, with the percentage of quartets in gene trees that agree with the ML topology (normalized quartet support) q1 (f1) being 0.62 (1989); higher than nodes supporting other deep splits in *Heliconius* (Doris + Wallacei + Silvaniform + Melpomene clade, with the Wallacei + Silvaniform + Melpomene branch).

**Fig.1.**
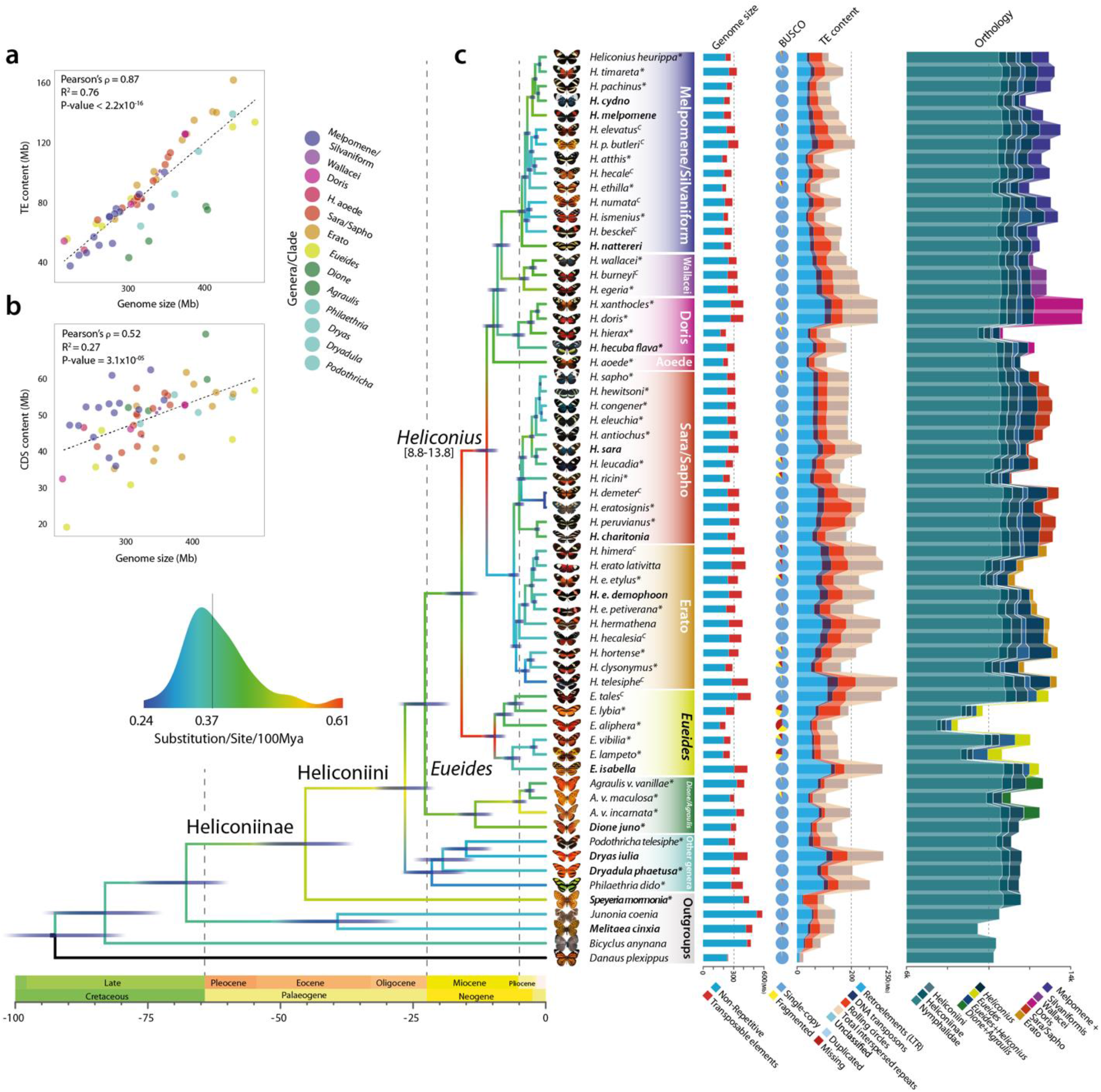
Phylogenetic, genomic, and proteomic comparisons among 63 Nymphalid butterfly species. **a, b** The contribution of transposable elements (TEs) and coding regions (CDS) to genome size variation across Heliconiinae, respectively. **c** From left to right: *i*) the dated species phylogeny built from the concatenated single-copy orthologous groups (scOGs) from all sequenced Heliconiinae and outgroups, using a combination of Maximum Likelihood and Bayesian Inference. The branch colour represents the number of substitutions per site per 100 Mya of that specific branch. Species names in bold indicate the species with chromosome- or sub-chromosome-level assemblies, asterisks indicate genomes assembled in this study, ^C^ curated assemblies; *ii*) genome assembly size, in red the TE fractions; *iii*) BUSCO profiles for each species. Blue indicates the fraction of complete single-copy genes; *iv*) bar plots show total gene counts partitioned according to their orthology profiles, from Nymphalids to lineage-restricted and clade-specific genes.

**Fig. 2.**
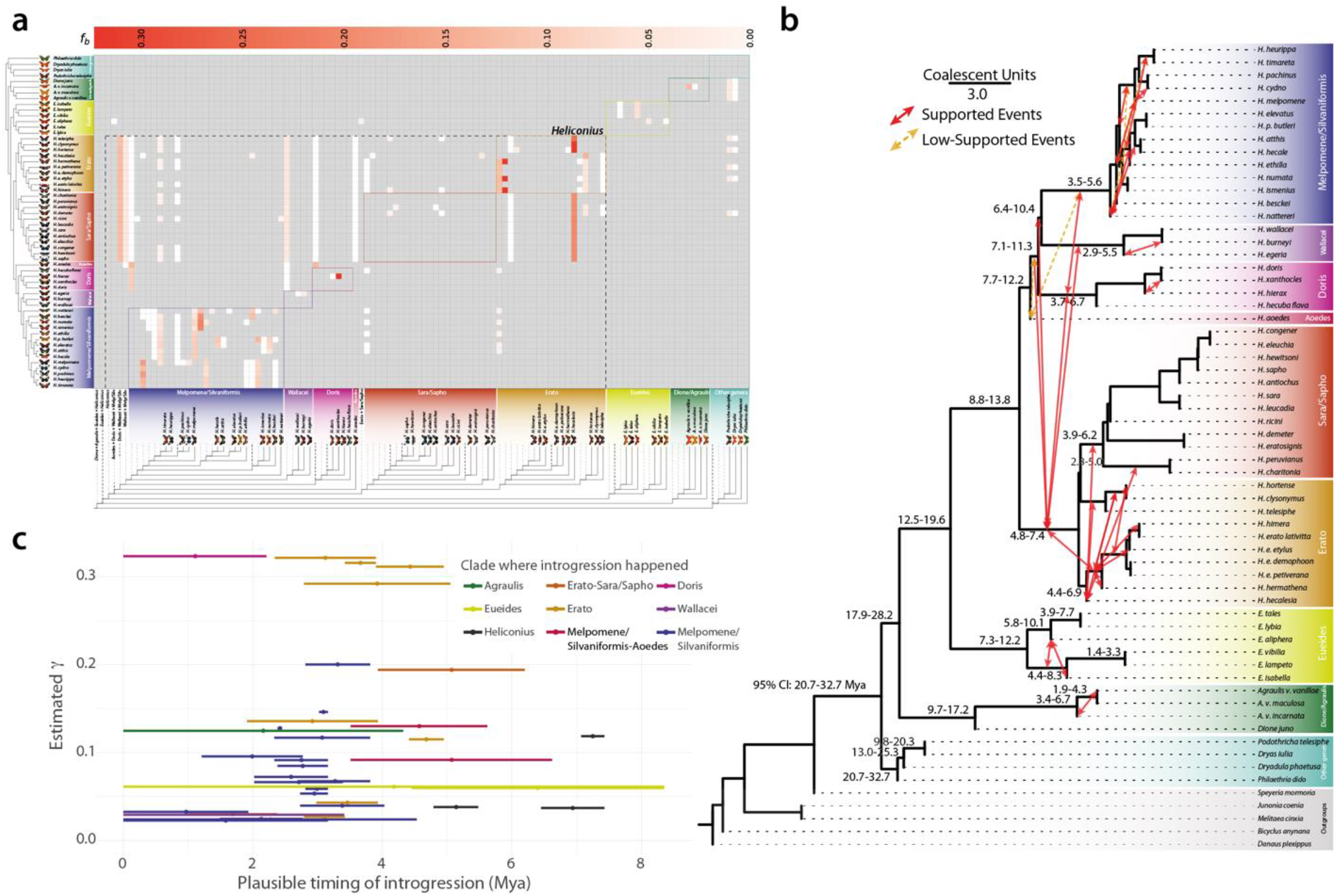
Patterns of introgression inferred for the Heliconiinae clade. **a** The matrix shows inferred introgression proportions as estimated from scOG gene trees in the introgressed species pairs, and then mapped to internal branches using the f-branch method. The expanded tree at the bottom of the matrix shows both the terminal and ancestral branches. **b** ASTRAL-III species tree derived from nucleotide gene trees, with mapped introgression events (red arrows) derived from the corresponding *f*-branch matrix. Yellow dashed arrows indicate introgression events with lower support (triplet support ratio < 10%). Branch lengths correspond to coalescent units. Numbers on nodes correspond to the confidence interval of the dated phylogeny (Fig. 1e). Note how, not only, most of the introgression events happen within clades and among time overlapped nodes, but also how the majority of introgressive events are affecting lineages with low CUs, indicating a lower barrier to gene flow. There seems to be only one introgression for H. aoede, which happened with the Silvaniform/Melpomene basal branch. Times inferred from the dating analysis summarized in Fig. 1. **c** Each segment indicates the confidence interval of the estimated introgression event (triplet support ratio > 10%). The circles indicate the average date.

*Heliconius* have also become key taxa for exploring the impact of gene flow and hybridization on adaptive divergence^5,7,13^. We therefore revisited this topic with our extended taxonomical range, adopting two very recent methodological approaches: the discordant-count test (DCT) and the branch-length test (BLT). Both tests reveal a lack of gene flow between basal Heliconiini nodes and those at the base of the *Heliconius* radiation, including the *H. aoede* split, but do identify several introgression events within major clades of Heliconiini (Fig. 2, and Supplementary Table 4, 5). Note, the putative lack of introgression at the basal node of Heliconiini is unlikely to be simply explained by a lack power in the statistical methods used to detect introgression. The Heliconiini split is dated between 20 and 30 Mya, and the same methodology, applied to the *Drosophila* radiation^14^, has identified introgression events dated over 20 My, suggesting that in principle the methods applied should be able to find introgression in our phylogenetic framework. The greatest number of introgression events were detected within *Heliconius*, specifically between the most recent common ancestors (MRCAs) of Erato + Sara/Sapho clade, the Doris + Wallacei + Silvaniform + Melpomene clade, and within the Erato clade. Interestingly, the Sara/Sapho clade shows very low rates of introgression, potentially reflecting a stronger barrier to gene flow^15^ between species in this clade, where females mate only once (monoandry), and males often mate with females as they eclose from the pupae (referred to as pupal mating)^16^. Across all branches, the estimated fraction of introgressed genome mostly varies between 0.02 γ to 0.15 γ, with a peak around 0.30 within the Erato clade (range of average γ estimates = 0.023– 0.323). Most introgression events also occurred in a restricted time frame within the last 5 Mya (Fig. 2b), and no significant relationship was found between the midpoint estimate of the timing of introgression and the estimated γ (Fig. 2b), indicating that the fraction of a genome that is introgressed within *Heliconius* does not depend on the timing of those introgression events (see Supplementary Material for more details).

### The Origin of Major *Heliconius* Lineages and Pollen-feeding

Pollen-feeding is one of the most important key innovations within *Heliconius* radiation. So far, all phylogenetic reconstructions based on molecular data^5,8^ place the non-pollen feeding clade Aoede (members of the genus formerly known as *Neruda*) within the *Heliconius* clade, suggesting a secondary loss in this lineage. The comparison of this lineage, represented in our data by *H. aoede*, with the pollen-feeding *Heliconius* species offers the intriguing possibility to understand the genetic basis of the traits related to pollen-feeding and potentially to solve the puzzle about its emergence. Specifically, we can test whether *i*) pollen-feeding emerged once, at the stem of *Heliconius*, with the Aoede clade outside *Heliconius s*.*s*.; *ii*) or if it emerged once with Aoede falling within *Heliconius*, and was secondarily lost in the Aoede clade; or *iii*) if it evolved independently in the Erato and Melpomene clades, with Aoede falling within *Heliconius* but without invoking trait loss. Using extensive genomic data in the form of scOGs, our data support the monophyletic status of the pollen-feeding *Heliconius* + *H. aoede* (Fig. 1c, and Supplementary Fig. 19). Specifically, *H. aoede* seems to cluster sister to the stem of three other clades: Melpomene/Silvaniform, Wallacei and Doris (Fig 1c, 2b). This position is strongly supported by bootstrap and concordance values (Fig. 1c and Supplementary Fig. 19, 20). A further assessment of nodal support was performed using the Quartet Concordance (QC), Quartet Differential (QD) scores, and Quartet Informativeness (QI) (within Quartet Sampling) to identify quartet-tree/species-tree discordance (see Methods). The position of *H. aoede* remained supported, with a strong majority of quartets supporting the focal branch (QC = 0.9), with a low skew in discordant frequencies (QD = 0) indicating that no alternative history is favoured, no signal of introgression is detected (i.e., QD < 1 but >0), and a QI of 1 indicates that the quartets passed the likelihood cut-off in 100% of the cases (Supplementary Figs. 22, 23; QC = 0.9). Leveraging the 63-way whole genome alignment and using *E. isabella* as reference, we further tested the robustness of this topology by inferring the local topology history across the 63 species. We generate non-overlapping windows of 10kb across the whole 63-way whole genome alignment and use them to infer ML trees at each window, exploring the frequency of possible topologies, the effect of introgression and incomplete lineage sorting (ILS) with a coalescent based method. From more than 43k non-overlapping sliding windows, ∼30k returned one of five main topologies. With the only purpose of exploring the monophyly of the pollen-feeding trait, we classified the topologies based on the position of *H. aoede* relative to the other *Heliconius* clades, *Eueides* and other non-*Heliconius* species (Supplementary Fig. 24a). The most frequent/supported topology (Topology 1, 49% of trees), shows the same relationships of our species tree reconstruction. Less frequent topologies (Topology 2,4, and 5, total of 16% of trees) also show *H. aoede* nested within *Heliconius*, while Topology 3, the topology that places *H. aoede* outside *Heliconius s*.*s*., has a frequency of 3.7% (see Supplementary Results and Supplementary Fig. 24). Aware of the possible impact of introgression and ILS on topology inference, we used the same non-overlapping sliding windows to infer the impact of those on different chromosomes, expecting Z chromosome to be less affected by both^7^. The fraction of the genome that introgressed (average *f*-branch statistic) across all triplet comparisons and coalescent units (CUs), as a proxy of ILS, from each chromosome and the Z chromosome versus all autosomes (see Supplementary Results and Supplementary Fig. 25, 26) show that, indeed, the Z chromosome has a lesser degree of introgression and ILS overall, but this effect does not change the topologies’ frequencies in favour of Topology 3, which stays ∼5% of the entire chromosome, whereas Topology 1 increase to 56%.

Overall, given the methods currently available for large phylogenomic datasets such as ours, the landscape of local history seems to confirm the species tree as the most consistent topology, with *H. aoede* clustering within *Heliconius* clades. This would likely exclude a single gain of pollen-feeding with no loss (*i*), leaving two nominally equally parsimonious scenarios; one gain at the stem of *Heliconius* clade, followed by one loss at the branch of *H. aoede*, or two independent gains at the base of Sara-Sapho/Erato and Doris/Wallacei/Melpomene/Silvaniform. For our purposes, this provides a hypothesis testing framework where, under the first scenario which is traditionally seen as most likely^10^, signatures of molecular innovation relating to the suite of traits linked to pollen-feeding may be expected to occur on the stem *Heliconius* branches, while the pattern of evolution on *H. aoede* is predicted to be linked to trait loss. In what follows, we use our phylogenetic framework to explore the evolution of genomic size, content and patterns of selection in key points of the Heliconiini radiation.

### Evolution of Genome Size and Content

Variation in genome size can be formalised in an “accordion” model^17^ where genomes gain, lose, or maintain its size in equilibrium in each species, due to a balance between expansions in transposable elements (TEs) and large segmental deletions. By reconstructing ancestral genome sizes at each node in the Heliconiini phylogeny, we found that the MRCA of Heliconiinae experienced a 30% contraction from ∼406Mb at stem of Heliconiinae to ∼282Mb for Melpomene/Silvaniform clade; while, at the same time, other branches leading to *Philaethria, Dryadula, Dryas* and *Podothricha*, and the Erato, Doris and Wallacei clades within the genus *Heliconius*, had independent expansions. Strikingly, *H. aoede* shows a loss of about 68 Mb, a fifth of its genome size (∼22%) from its ancestral node (Fig. 3).

**Fig. 3.**
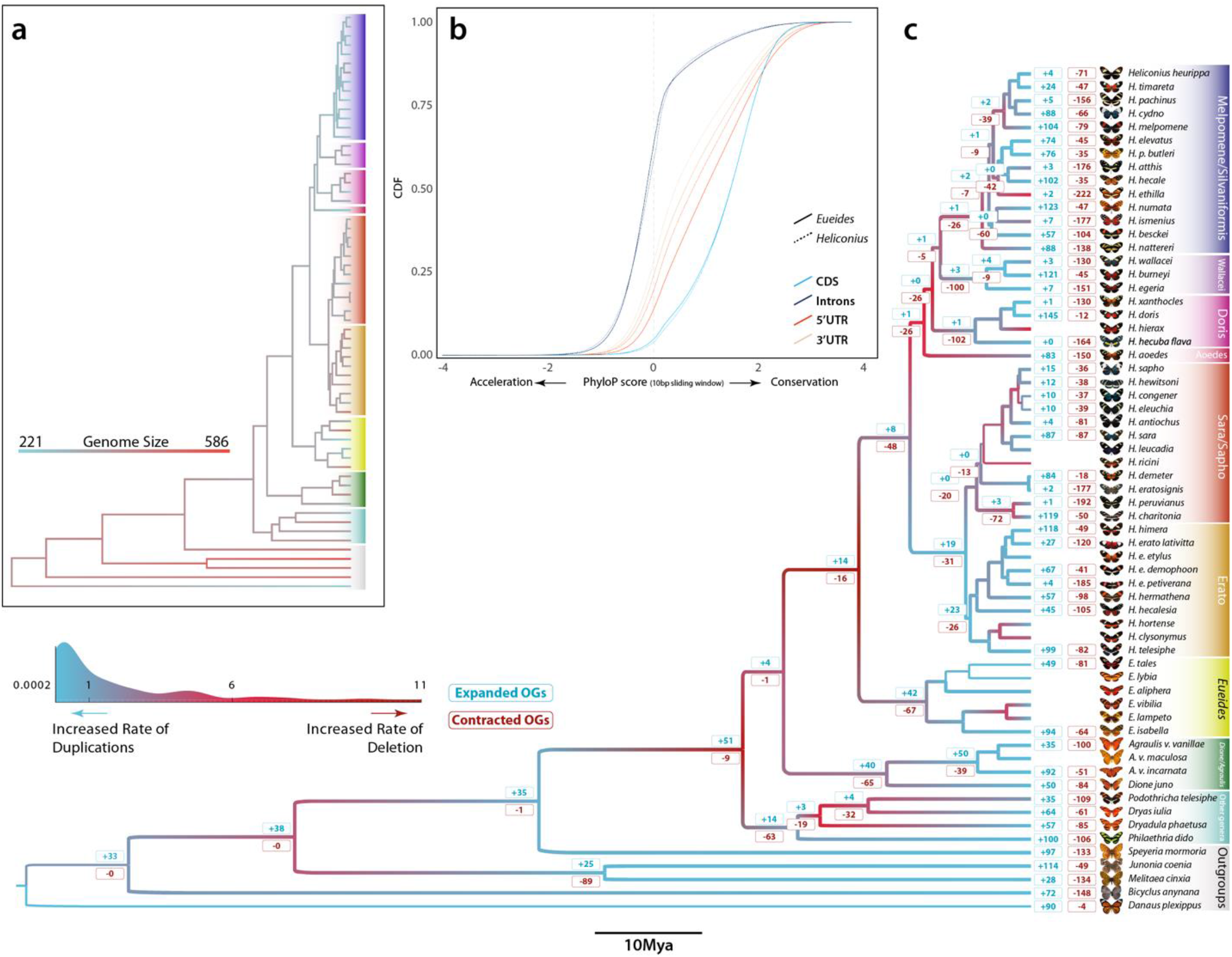
Genomic dynamics, acceleration/conservation rates and ortholog copy number evolution. **a** Ancestral genome size reconstruction of Nymphalids inferred by ML approach. **b** Cumulative distribution functions (CDFs) of scores for selected annotation classes (CDS, introns, 5’ and 3’ UTRs) as computed by the subtree scores for *Eueides* and *Heliconius* clades. PhyloP scores at sites of different annotation classes, based on the LRT method and multiple whole 63-species genome alignment. Positive scores indicate conservation, and negative scores indicate acceleration (CONACC mode) in a 10-bp sliding window. **c** Branch colours indicate the ratio between the rate of duplications (duplicated Mb/Mya per branch) and deletions (deleted Mb/Mya per branch) across the Nymphalid phylogeny. In general, red shifts indicate an increased rate of deletion over rate of duplication; the opposite is true for blue shifts. Numbers at nodes correspond to the amount of expanded (blue) and contracted (red) ortholog groups (values are shown for main branches and most complete genomes, see Methods).

There is a remarkable difference is species richness between the two sister genera, *Heliconius* and *Eueides*, but both seemed to have experienced an accelerated rate of substitution (Fig. 1c). We explicitly tested which genomic compartments (CDS, introns, 5’-UTR, and 3’-UTR) contribute to the change in substitution rate. We did so by calculating CONACC scores and assessing departures from neutrality (Fig. 3b). Between the two genera, we identified an enrichment for higher CONACC scores in *Heliconius* for CDS and introns, compared to the same compartments in *Eueides*, a trend that is inverted for the two UTR regions (Wilcoxon rank-sum test ‘two-sides’ *P* value < 2.2×10^−16^). This suggests an increased tendency for clade-specific selection, also confirmed by the fast-unconstrained Bayesian approximation method (FUBAR), which showed that *Heliconius* have more sites under purifying selection and positive selection per codon compared with *Euiedes* (Supplementary Fig. 27). *Heliconius* has 2.5x more sites under purifying selection per codon than *Eueides*, suggesting that the higher CONACC scores in *Heliconius* are likely due to higher degrees of purifying selection.

We next explored the relationship between transposable elements (TEs) and genome size, and their effect on gene architecture. We found that larger genomes tend to have a distribution of intron length skewed towards longer introns (Supplementary Fig. 18a), with a positive correlation between median intron length and total TE content (Supplementary Fig. 28; Pearson’s *ρ*=0.72; R^2^=0.51). Long introns also accumulate significantly more TEs then expected by their size (Supplementary Fig. 29; Wilcoxon rank-sum test *P* value = 2.13×10^−13^), with the effect of changing gene structure more than gene density (Supplementary Fig. 30). This suggests that selection may have favoured a reduction in TEs in intergenic regions, perhaps to avoid the disruption of regulatory elements^18^, consistent with TEs largely accumulating in the tails of chromosomes^19^. Although intron size varies significantly, the rate of gain/loss of introns, and the intron retention from the MRCA of Nymphalids, shows a relatively stable dynamic over the last 50 Mya in Heliconiinae, with no significant shift among species, and ∼7% of ancestral intron sites retained across species (Fig. 4c). A similar pattern was reported in *Bombus*^20^, but our results differ from drosophilids and anophelines, which show significantly higher intron turnover rates^21^.

**Fig. 4.**
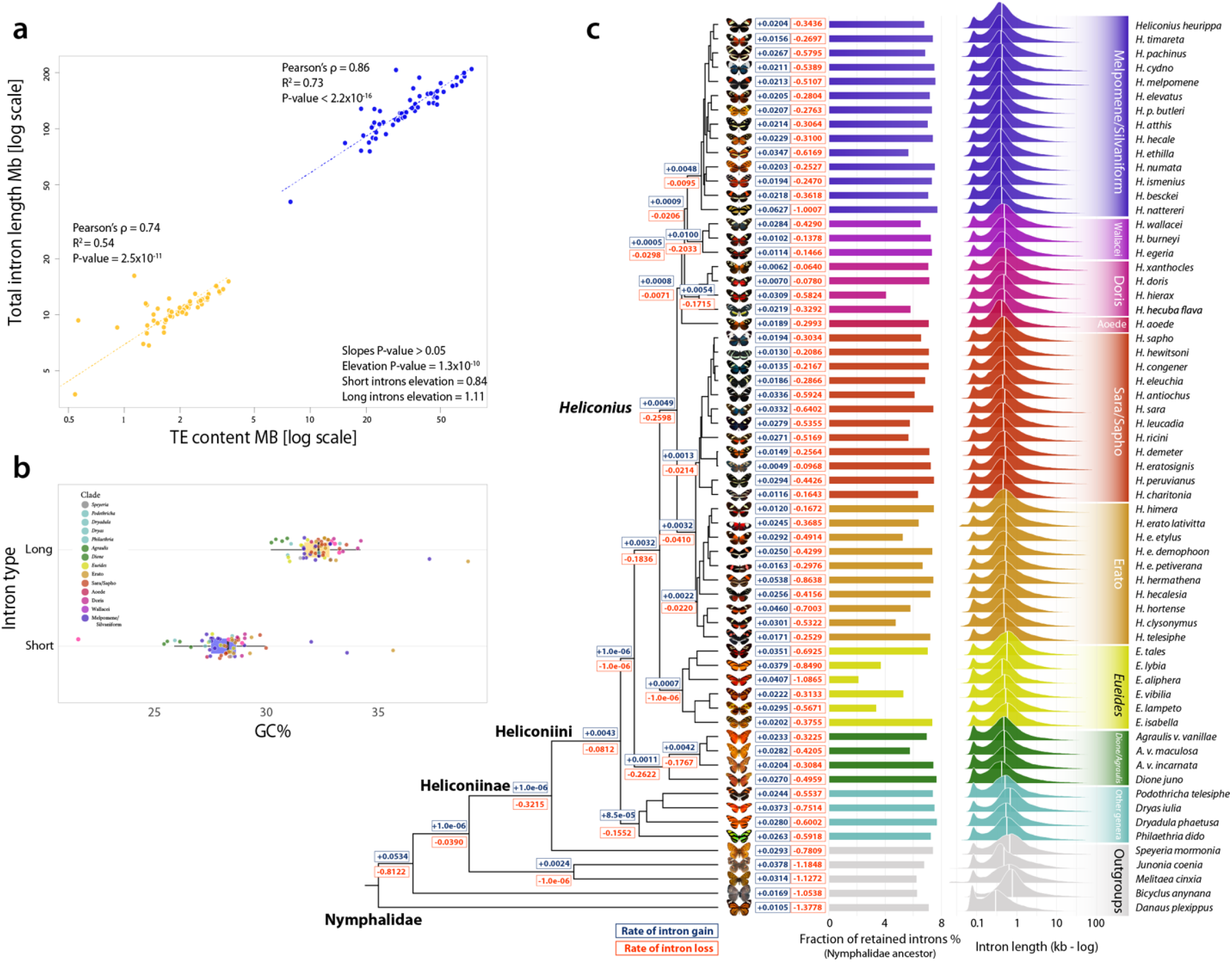
Composition and intronic evolution. **a** Log-log scatter plot showing high correlation between intron length and Transposable Element (TE) abundance. The significantly higher elevation of longer introns compared with short introns, indicates that large introns are more affected by TEs; short intron elevation = 0.84; long intron elevation = 1.11; *P* value = 1.3×10^−10^. **b** Box plot of GC content and intron size (Wilcoxon rank-sum test *P* value < 2.2×10^−16^). **c** Rates of intron gain (blue) and loss (red) across Nymphalid phylogeny, and fraction of retained introns from the Nymphalid ancestor. On the far left, the log scale distributions of intron lengths, white vertical bars indicate the median values.

### Expansion and Contraction of Gene Content

The Heliconiini tribe shows a diversity of key innovations in different aspects of their physiology and adaptation. These aposematic butterflies *de novo* biosynthesize their toxins when similar compounds are not available from their obligatory larval hostplant (Passifloraceae) for sequestration – a process called biochemical plasticity. These toxins not only make heliconiines distasteful to predators, but also play important role during mating^22,23^. The Heliconiini also produce complex and diverse bouquets of pheromones^24^, which can play an important role in speciation through the formation of pre-zygotic reproductive barriers, ultimately reducing gene flow and facilitating speciation. *Heliconius*, however, specifically show other traits, including an extended lifespan and increased neural investment^25^ compared with other butterflies and sister clades, which are thought to have evolved alongside pollen-feeding^10^. We tested if the origin of these suites of traits are associated with gene expiations/contraction at key points in the phylogeny, modelling the turnover rate of ortholog group (OG) size with CAFE v5^26^ for 10,361 OGs using the 52 most complete genomes (BUSCO score ≥ 90%). The analysis identified 656 OGs that vary significantly in size across the phylogeny. The estimated gene turnover (λ) was of 0.006/gene gain-loss/Mya. This is relatively high compared with rates for *Bombus* (λ = 0.004) and anopheline species (λ = 0.003)^20,21^, but similar to drosophilids (λ = 0.006)^27^. The base of the phylogeny showed relatively strong OG expansions, with few contractions, followed by stasis. While *Dione* + *Agraulis* and *Eueides* stems have similar proportions of expanded/contracted OGs, *Heliconius* shows 48 contracted OGs but only eight expanded OGs (Fig. 3c, Supplementary table 7) suggesting the phenotypic innovations that occurred in this branch were not due to widespread gene duplication.

Several OGs were identified to be expanded multiple times across the phylogeny and some of these may be directly associated to previously described key innovations/phenotypic traits across Heliconiini. For example, we find that cytochrome P450 (P450s) genes expanded in the common ancestor of the subfamily Heliconiinae, the tribe Heliconiini, the *Dione* + *Agraulis* stem, and within the genus *Heliconius* in the Erato group and Silvaniform*/*Melpomene stems. In insects, P450s play important roles in the detoxification of specialized metabolites, hormone biosynthesis/signalling, and biosynthesis of cyanogenic glucosides in heliconiine butterflies, which form the basis of their chemical defence^23^. Notably, a range of diet related OGs are also highlighted: Glucose transporters and Trypsins expanded several times in Heliconiinae, Heliconiini, *Eueides*, and the Silvaniform/Melpomene stem. Although glucose transporters play an important role in energetic metabolism in all animals, in phytophagous insects they are also hypothesised to be involved in the sequestration and detoxification of specialized metabolites from plants^28^. There are also expansions in Lipases enzymes, and OGs linked to in energetic metabolism and in pheromone biosynthesis in the Sara/Sapho + Erato stem and Silvaniform/Melpomene clades. At the stem *Heliconius* species there is one duplication of *methuselah*-like, a G-protein coupled receptor, involved in oxidative stress response, metabolic regulation, and lifespan^29^, together with *Esterase P*, and a *juvenile hormone acid methyltransferase* (*jhamt*) which expanded three times. Taken together these expansions events offer good candidates for pathways which may be linked to the derived life history traits and chemical ecology of the Heliconiini.

We further expanded the previous unsupervised analysis by focusing on 57 gene families (GF), which includes a range of biological functions (Supplementary Table 8). We used measures of “phylogenetic instability” and the gene turnover rate (λ, CAFE), to explore their dynamics. The average instability score was 37.45 while the average λ is 0.005, with the number of OGs per family positively correlated with λ (Pearson’s *ρ* = 0.42). Sodium/calcium exchanger proteins and the Hemocyanin superfamily show the highest instability and turnover rates (Supplementary Fig. 39). This analysis also identifies GFs which expanded in key periods of heliconiine diversification, including Hemocyanins, Lipases, Trypsins and Sugar transporters and the Major Facilitator Superfamily (Fig. 5a). The most notable are the P450 CYP303A1-like gene (Supplementary Fig. 40), a highly conserved protein in insects that has a pivotal role in embryonic development and adult moulting^30^, and two Hemocyanins, a hexamerin storage homolog expanded in the *Dione* + *Agraulis* + *Eueides* + *Heliconius* clade, and the arylphorin homolog, expanded in *Eueides* and *Heliconius*. These hexamerins function as storage proteins, providing amino acids and energy for non-feeding periods, such as molting and pupation, and may also transport hormones^31^. We further characterised OGs within each GF, aiming to test for correlated gene expansions/contractions and shifts in selective pressures between *Eueides* and *Heliconius* (see Supplementary Results for more details), and found that Hemocyanins show, among several GFs, evidence of divergent selection regimes (*ω*) between *Eueides* and *Heliconius*, alongside Trypsins, Protein kinases, P450s, Sugar transporters, Ion and ABC transporters (Fig. 5b). Curiously, a contraction in the Hemocyanin superfamily was only observed in *H. aoede*, in our data, the only *Heliconius* species that do not to feed on pollen, marking hexamerins a potential mechanistic link to the divergent strategies for nitrogen storage in pollen-feeding *Heliconius* (Fig. 5a). However, our analyses indicate that a number of putatively important changes in gene family size not only occurred at the stem of *Heliconius*, but also in more basal branches at level of subfamily and tribe, before the adaptive radiation of *Heliconius*.

**Fig. 5.**
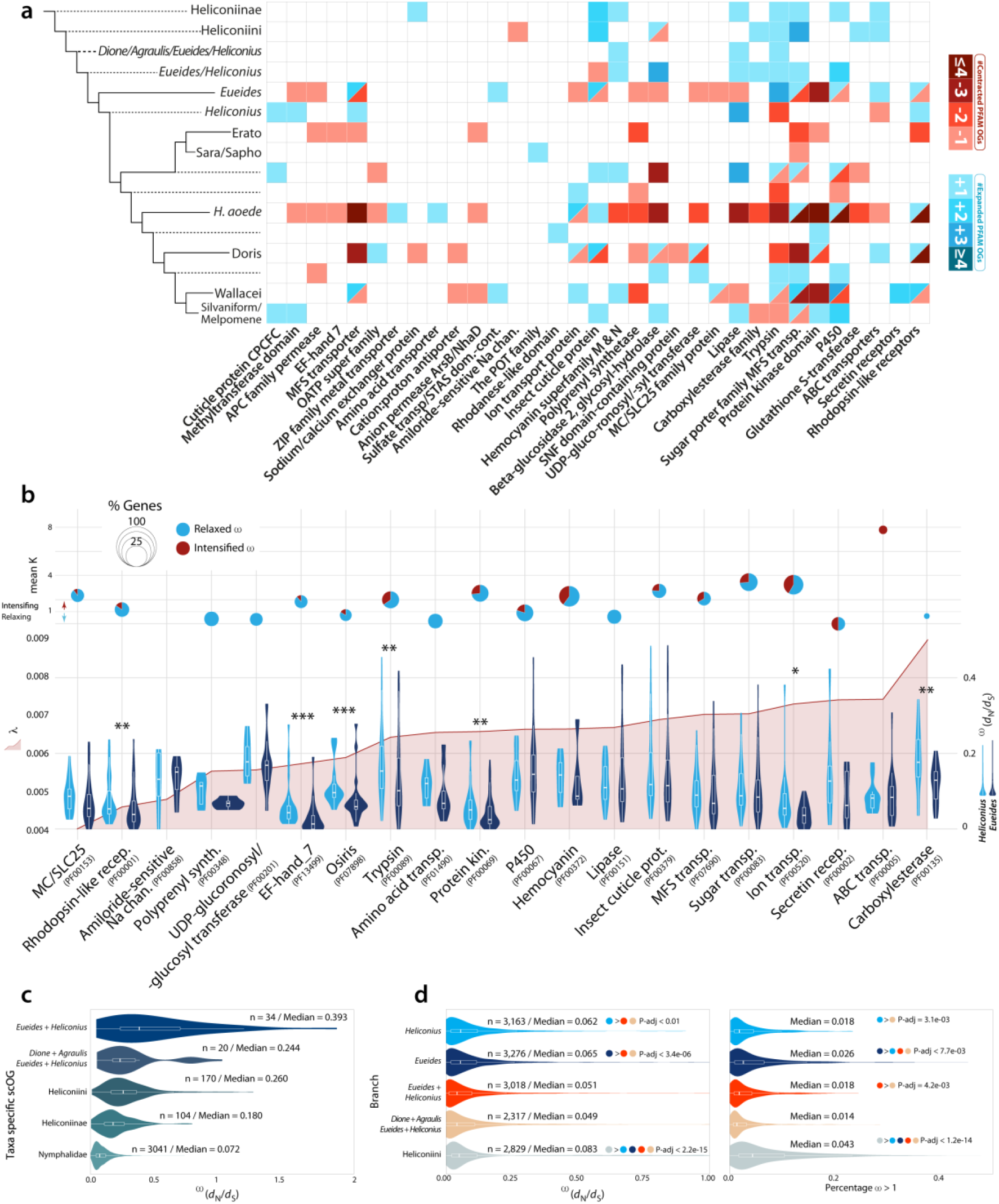
Differential evolutionary rates gene families and scOGs across Heliconiini butterflies. **a** Heatmap showing the different expansions and contractions in multiple gene families. Several gene families have been contracted in *H. aoede*. **b** Plots showing different evolutionary features of some of the analysed gene families (minimum of 3 genes in *Eueides spp*. and Heliconius *spp*.). At the top section, dynamic pie charts showing mean K value (selection intensifier parameter). Values below one indicates a relaxation, while above one indicates intensification towards diversifying positive selection. The size of the pie charts indicates the fraction of genes under intensification (red) and relaxation (blue), and it is scaled according to the proportion of genes for which K was significantly different from *H*_0_ (No difference) (see Methods). For different gene families the panel below shows the gene turnover rate (λ) (left y-axis); right y-axis shows the distributions of mean ω for near-scOGs (see Methods) in Eueides and Heliconius (right y-axis). Asterisks indicate significant shifts between Eueides and Heliconius (Wilcoxon rank-sum tests). **c** Violin plot showing the distributions of mean *ω* rates (*d*_N_/*d*_S_) in scOGs according to their lineage-specificity. **d** Distribution of mean ω rates (left) for scOGs on six branches of Heliconiini, and the proportion of genes for which ω is higher than one (right).

### Selection Across the Heliconiini Radiation

Selection regimes shaping Heliconiinae coding genes were further investigated using the adaptive branch-site random effects likelihood (aBSREL) method. Again, we aimed to examine positive selection at the *Heliconius* stem, and contextualise these patterns by testing and measuring the degree of diversifying positive selection at more basal branches. First, when single-copy orthologous groups (scOGs) are classified according to their phylogenetic attribution (*i*.*e*., where they appeared throughout the phylogeny), they show a trend towards increased purifying selection from young to older genes (Fig. 5c), suggesting that genes become more stable with time, probably reflecting increased functional importance. The signature of diversifying positive selection was assessed on five basal branches of the Heliconiinae phylogeny where key ecological transitions occur. From the Heliconiini to *Dione + Agraulis + Eueides + Heliconius, Eueides + Heliconius, Eueides*, to the *Heliconius* stem. Overall, the Heliconiini branch evolved under the strongest selection, followed by the *Eueides* and *Heliconius* branches, and finally by the *Eueides* + *Heliconius* branch (Fig. 5d). The number of genes with a signal of diversifying positive selection varies between branches, with the *Dione + Agraulis + Eueides + Heliconius* and *Heliconius* stems having the highest number of enriched biological processes (BPs), followed by Heliconiini and *Eueides* stem, and *Eueides + Heliconius*. A notably high proportion of branches are enriched for BPs relating to neuronal development and cellular functions, including the regulation of hippo signalling, stem cell differentiation and cell-cell adhesion, and genes associated with asymmetric division (Supplementary Tables 9-11). Using a network-based approach, which integrates both primary and predicted interactions to predict gene function, we examined connections between selected genes. Although the amount of network interactions shows a significant degree of connectivity (absolute number of interactions) in the branches leading to *Dione + Agraulis + Eueides + Heliconius* (834 interactions), *Eueides + Heliconius* (627), *Eueides* (531), Heliconiini (410), and *Heliconius* (320), the network density shows a different picture, with *Heliconius* having a markedly higher density (the portion of the potential connections in a network that are actual connections) with ∼0.3 versus ∼0.2 for the other networks, in the case of BP networks (Supplementary Fig. 41,Supplementary Fig. 42, and Supplementary Table 11). The enriched molecular functions (MFs) in this densely connected *Heliconius* network are characterised by BPs related to response to DNA damage/repair, neuroblast division, and neural precursor cell proliferation, glial cell development, cell-cell junction assembly, asymmetric stem cell division. This concentration of neurogenesis-related functions differs from enrichment in other networks, which appear more variable. Finally, we note multiple genes that show a signature of diversifying positive selection on more than one branch. One of them is the *Notch* homolog, an essential signalling protein with major roles in developmental processes of the central and peripheral nervous system^32^. Notch regulates neuroblast self-renewal, identity and proliferation in larval brains, and is involved in the maintenance of type II neuroblast self-renewal and identity^33^. Overall, these findings support the idea that many genomic changes that can be putatively linked to key *Heliconius* traits reflect a continuation or exaggeration of changes that occur in earlier Heliconiini lineages, suggesting a more gradual pattern of genetic evolution that precedes the adaptive radiation of *Heliconius*.

### Acceleration of Conserved Non-Exonic Elements (CNEEs)

The scan for diversifying positive selection on protein-coding genes showed interesting patterns that could be correlated to the evolution of phenotypic traits in Heliconiini. However, as we have seen, selection on the stem of *Heliconius*, although strong, does not seem to affect a high number of genes. We therefore expanded our scope to non-coding regions, specifically to regions of the genome that are conserved across the phylogeny but show altered patterns of evolution on the *Heliconius* stem. Comparative genomics approaches have assumed a fundamental role in the identification of conserved and functionally important non-coding genomic regions^34–37^. One of the most prominent hypotheses is that these regions function as *cis*-regulatory elements (such as enhancers, repressors, and insulators) and determine tissue-specific transcripts during developmental stages. To determine the extent of non-coding molecular evolution on the radiation of *Heliconius* butterflies, we compiled a total of 839k conserved elements (CEs) across the 63-way genome alignment (for a comparison, 1.95M CEs were found in birds^38^), leveraging a statistically neutral substitutional model, which considers phylogenetic distances and species relationships, to provide a more rigorous measure of actual evolutionary constraint^39^. Of the total CEs, 473k (56%) overlap with protein coding loci and 143k (30%) with coding exons, with 680k classified as conserved non-exonic elements (CNEEs), which were subsequently filtered (see Methods) to obtain a final set of 430,606 candidate CNEEs from the 63-way whole-genome alignment (811,696 in birds^38^); 202k intronic and 227k intergenic, for a total data set of 46,877,100 base pairs of aligned DNA.

We first checked for evidence of putative regulatory function by looking at the relationship between CNEEs and accessible chromatin, using ATAC (assay for transposase-accessible chromatin) peaks of 5^th^ instar caterpillars from two tissues, brain and wing imaginal disc^40^. We found that in both tissues CNEEs overlap ATAC peaks twice as often as expected under a random distribution (permutation *P*-value < 0.0001), with brain tissue having a slightly higher increase of 2.4 fold-enrichment, compared with the imaginal disc tissue (2.0 fold-enrichment). This is in spite of imaginal discs having twice as many ATAC peaks, covering twice the genomic region. This indicates that our annotated CNEEs are consistent with being putative functional elements and suggests that regulatory regions associated with brain tissue may be under more constraint, with a more conserved regulatory architecture.

Because of the putative regulatory relevance of CNEEs we applied a Bayesian method^41,42^ to detect changes in conservation of these elements at the stem of *Heliconius*, aiming to identify putative regulatory regions responsible for morphological and physiological adaptations of these butterflies. In total, we found that approximately half of the CNEEs (51%) experienced an acceleration in evolutionary rate at some point in the phylogeny. Around 95k elements experienced acceleration under a “full model” (M2), meaning that the latent conservation states **Z** (−1: missing, 0: neutral, 1: conserved, or 2: accelerated) can take any configuration across the phylogeny, while 122,445 elements best fit the lineage-specific model (accelerated on the *Heliconius* stem branch; M1), where substitution rates on the branches leading to target species are accelerated whereas all other branches must be in either the background or conserved state; of them 2,536 were accelerated (aCNEEs) at the stem of *Heliconius*. Among this list, we tested if there is enrichment of aCNEEs in accessible chromatin of brain and wing imaginal disc and found that in both tissues there was a similar fold-enrichment of 1.08 and 1.09, for brain and wing tissue, respectively (*P*-value = 0.04 for the brain tissue; *P*-value < 0.001 for imaginal disc). We then checked for enrichment of aCNEEs across genes, as well as their spatial distribution across the genome to identify genes most affected by the acceleration, or large regulatory hubs. We found 37 genes that harbour more aCNEEs in their putative regulatory domains than expected by chance (Supplementary Table 12). Among them, there are multiple genes linked to axon pathfinding^43,44^ (two genes homologous to *Uncoordinated 115a, Unc-115a, Eisa2300G23*: 5 aCNEEs; *Eisa2300G24*: 6 aCNEEs; adj. *P*-value < 0.026; *Multiplexin, Mp, Eisa1200G485*: 5 aCNEEs; adj. *P*-value < 0.026), synaptic pruning and transmission^45^, and long term memory^46^ (*Beaten path* Ia, beat-Ia, *Eisa2300G476*: 3 aCNEE; adj. *P*-value = 0.022; *Tomosyn, Eisa1400G28*: 4 aCNEEs; adj. *P*-value = 0.026). We also find examples such as *Nicastrin* (*nct*), which encodes a transmembrane protein and a ligand for Notch (N) receptor (*Eisa0300G576*: 3 aCNEEs; adj. *P*-value = 0.0014), and is required for neuronal survival during aging and normal lifespan, functioning together with a *Presenilin*-homolog (*Psn*)^47^ (*Eisa1800G396*: 1 aCNEE) which, although not enriched, also has one aCNEE in its regulatory domain. Finally, two pheromone binding proteins (*PhBPloc02ABP1*: 2 aCNEEs; adj. *P*-value = 0.026; *PhBPloc08ABPX*: 3 aCNEEs; adj. *P*-value < 0.05) and a sugar taste gustatory receptor (*Eisa0300G244*: 3 aCNEEs; adj. *P*-value = 0.041) are also highlighted as having multiple aCNEEs in their regulatory domain on the stem *Heliconius* branch.

The spatial enrichment analysis also highlighted 55 genomic regions significantly enriched upon *P*-value correction (Supplementary Table 13). Two of these correspond to a 150 kb (8 aCNEEs) and 120 kb (4 aCNEEs) gene deserts, meaning they contain no annotated protein coding gene. In proximity of these regions are mainly coding transposable elements, or viral ORFs, such as x-elements or retrovirus-related Pol polyproteins of *Drosophila*, and nearby *collagen alpha-1(III) chain-like, Argonaute 2, Osiris 21* (*osi21*) and *spalt major* (*salm; Eisa0200G420*: 8 aCNEEs), an important zinc finger transcriptional repressor that mediates most decapentaplegic (dpp) functions during the development of the wings. The product of *salm* is also required for cell specification during the development of the nervous system, muscle, eye or trachea^48^. Together with the notion that gene deserts have pivotal regulatory functions^49^, this makes these two regions important candidate regulatory hub for developmental processes in *Heliconius*. A further enriched genomic region is located on chromosome 20 (Fig. 6). This region harbours eight aCNEEs distributed across two putative regulatory domains of two genes, both homologs of *osa*, which encodes for a subunit of the Brahma-associated protein (BAP) chromatin remodeling complex, part of the SWI/SNF chromatin-remodeling complexes. This complex functions to alter the accessibility of transcription factors to genomic loci. As such, it plays important gene regulatory roles in multiple contexts^50^. In *Drosophila*, it controls escorting cell characteristics and germline lineage differentiation^51^, but the complex is also implicated in inducing the transcription of *crumbs* (*crb*), which we also found to have one aCNEEs in its putative regulatory domain. Crumbs, in turn, is a transmembrane protein which negatively regulates the Hippo signalling cascade, and plays an integral role in cell proliferation and tissue growth regulation^50^. Additionally, the silencing (by RNAi) of different subunits of the BAP complex results in disrupted short- and long-term memory, while direct silencing of *osa* impaired the retention of long-term memory^52^. Given that long-term memory is thought to be stable across longer periods in *Heliconius* than related genera^25^, these reflect clear candidate loci of interest. We also examined evidence of GO term functional enrichment among the 2,536 *Heliconius*-specific aCNEEs (Supplementary Fig. 43-45), using different approaches which resulted in similar enriched categories (Supplementary Table 14). Specifically, 36 aCNEEs are linked to strongly enriched transcription factors and receptors related to imaginal disc-derived wing morphogenesis (*e*.*g*.: *dl, osa, ser, lgs, dll, fz2, sfl*), and retinal cell differentiation (*e*.*g*.: *salm, emc*), 36 aCNEEs near 14 genes are related to the Notch signaling pathway (*e*.*g*.: *agxt, ham, got1, nct, psn, noc, wry, nedd4*), and 20 aCNEEs near 11 genes are related to feeding behaviour (*e*.*g*.: *for, 5-ht2a, dip-kappa*).

**Fig. 6.**
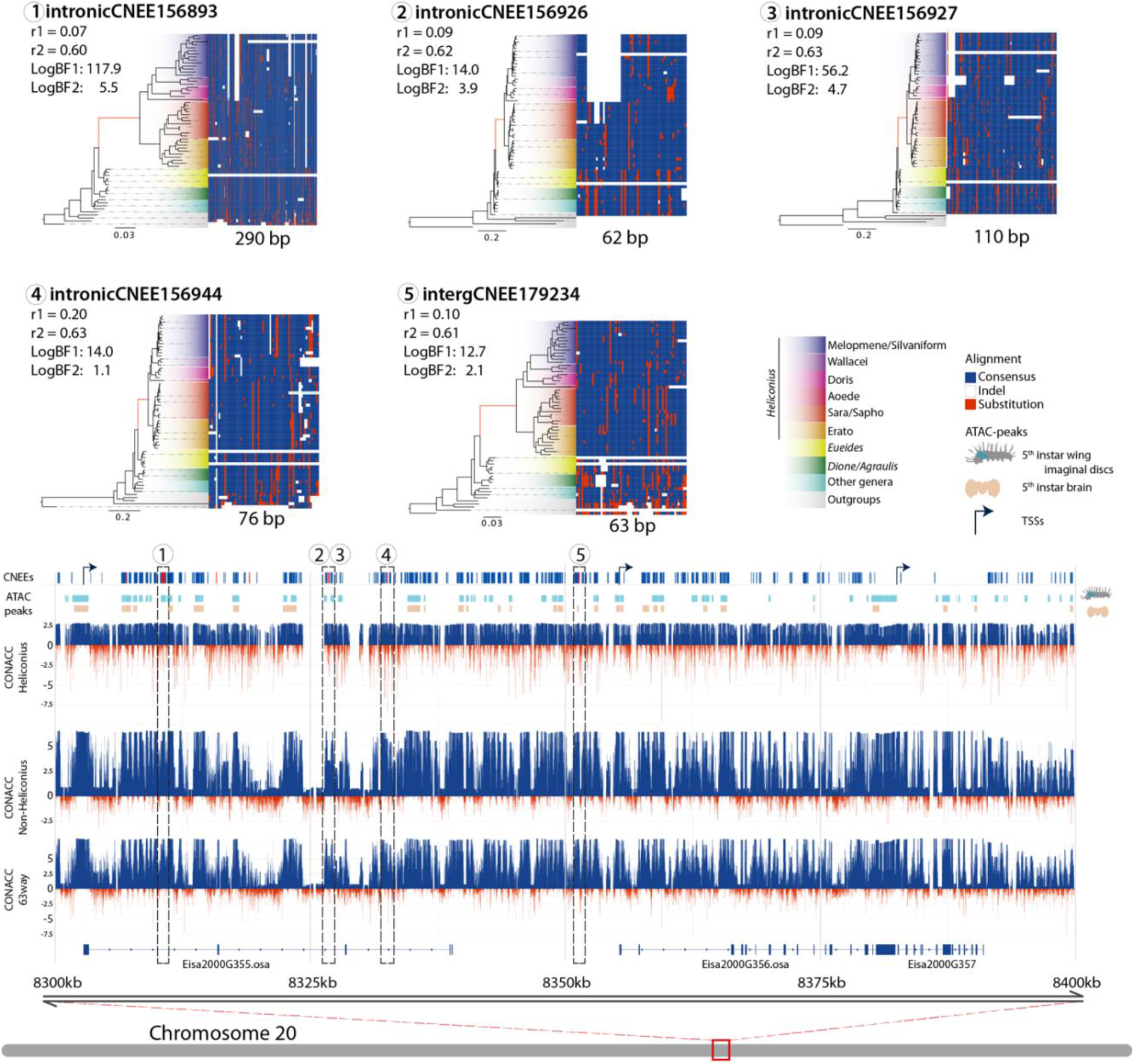
Chromosome 20 enriched genomic window. Diagram showing the distribution of CNEEs in one of 100 kb enriched window across the reference genome of *E. isabella*. From the bottom-up, the figure shows three genes, two of them homologs of the *Drosophila osa*. Above that are the CONACC scores obtained from the full alignment (63 species), for only the non-*Heliconius* species, and only the *Heliconius* species. In red the negative values indicate the acceleration of a given position of the alignment, and in blue the positive values indicate conservation. Above that are the ATAC peak distributions of two tissues from 5^th^ larva instar, brain (in brown) and imaginal disc (in aquamarine), shown alongside the distribution of CNEEs in the region in dark blue with the eight aCNEEs. Numbers indicate the five aCNEEs selected as examples of the *Heliconius* aCNEEs, which are expanded at the top of the figure. In these examples, the alignments show conserved (nucleotides similar to the consensus, in blue) and accelerated sequences (nucleotides that differ from the consensus, in red). For each of the five aCNEEs the species phylogeny of the Nymphalids is shown where the branch lengths indicate the acceleration of the evolutionary rate for each given aCNEE. The branch that corresponds to the *Heliconius* stem is in red. For each aCNEE the two log-BFs and conserved (*r*_1_) and accelerated rate (*r*_2_).

### Candidate Genes for Derived Traits of *Heliconius*

Within the Heliconiinae, *Heliconius* display a number of divergent traits and innovations^10^. Here, we highlight how our results reveal new biological insights into these traits, focusing on two case studies; changes in neural composition in Heliconiini, and the enzymatic processes associated with breaking down pollen walls to aid their digestion during pollen feeding. These two examples, illustrate the potential of large, densely sampled genomic datasets to both generate and test adaptive gene-phenotype hypotheses, using both unguided and more targeted analyses.

Within in central brain, mushroom bodies are paired organs that receive visual and/or olfactory information, and play a pivotal role in learning and memory^53^. These structures show huge variation across Heliconiini, but a particularly large expansion occurred at the *Heliconius* stem, where mushroom body volume and neuron number more increased by several-fold, accompanied by a major shift towards increased dedication to processing visual information^11,54^. These changes are accompanied by enhanced learning and memory performance^25^, and likely facilitate the foraging strategies deployed during pollen feeding. However, the molecular mechanisms underpinning these events – or, indeed, any case of mushroom body, or brain expansion in insects - are unknown. Given the lack of variation in closely related species suitable for alternative approaches, comparative genomics reflects the best route to identifying genes linked to this shift in brain morphology. Our selection analyses highlight pathways that could regulate neural proliferation. These include the Hippo signalling pathway, which regulates cell growth and proliferation of neural stem cells and neuroblast quiescence^55^. Multiple and repeated signs of diversifying selection are identified on genes related to the Hippo signalling pathway, including *Focal adhesion kinase* (*Fak*), *lethal (2) giant larvae* (*lgl*), *Sarcolemma associated protein* (*Slmap*), and *Akt kinase* (*Akt*), on the *Dione + Agraulis + Eueides + Heliconius* stem, which regulate cell polarity, asymmetric division and cell proliferation^56^, and two other genes, *Moesin* (*Moe*) and *F-box and leucine-rich repeat protein 7* (*Fbxl7*), in the *Eueides + Heliconius* stem. Moe drives cortical remodelling of dividing neuroblasts^57^, while Fbxl7 affects Hippo signalling pathway activity^58^. Finally, *Ctr9, dachsous* (*ds*), *falafel (flfl)*, and locomotion defects (loco) were identified at the *Heliconius* stem (Supplementary Table 9). Ctr9 is involved in the proliferation and differentiation of the central nervous system^59^, Flfl is required for asymmetric division of neuroblasts, cell polarity and neurogenesis in mushroom bodies^60^, Ds is a cadherin that interacts with the Hippo signalling pathway^61^, and loco is an activator of glial cell fate, essential cells in efficiently operating nervous systems^62^. Similarly, our analysis of conserved non-coding elements reveals multiple loci nearby genes with known roles in neural development, synaptic pruning, and long-term memory. Collectively, these provide the first candidate loci linked to mushroom body expansion in any insect and provide ample gene-phenotype hypotheses for further investigation.

Despite being a keystone innovation in *Heliconius*, similarly little is known about the mechanism underpinning pollen-feeding itself. Saliva probably has an important role in the external, enzymatic digestion of the pollen wall^10^. The leading candidates for these enzymes are serine proteases, homologs of the silkworm cocoonase^63,64^, which digests the cocoon during eclosion^64^. Because butterflies do not produce a cocoon, it has been proposed that the duplications of cocoonase orthologs may have been co-opted to digest pollen^63^. Given our order of magnitude larger sample, and having not highlighted this gene family in our unguided analysis, we re-evaluated the evolution of these genes by reassessing the evolutionary history of this gene family, and evidence of gain-of-function. We identified 233 cocoonase loci (Supplementary Table 15) across all Heliconiinae and found that the duplications not only predate the split between *Heliconius* and *Eueides*, but affect the whole Heliconiini tribe and its outgroup *S. mormonia* (Fig. 7a). All species have at least four copies, located at the minus strand of chromosome 15 with remarkable conserved synteny (Fig. 7b and Supplementary Table 16). Substrate, cleavage, and active sites of the functional domain show very high conservation throughout the dataset. Three independent tandem duplications from the same original copy are very likely responsible of the emergence of *Coc1A, Coc1B, Coc2* and *Coc3* (Fig. 7a). High level of purifying selection is detected across the four OGs (Fig. 7a). A scan of all internal branches for signs of diversifying positive selection shows that the branches of *D. juno Coc2* in-paralogs; the stems of all *Heliconius Coc1A* and *Coc1B*, and the two branches of the Silvaniform/Melpomene *Coc2* out-paralog, show signs of positive selection. Two of these events involve loci from the non-pollen feeding *H. aoede*. We therefore tested if these loci show signs of relaxation in this species. Surprisingly, while no significant differences were detected for *Coc1B*, an intensification of selection was detected for *Coc1A* (*K* = 1.41; *P*-value = 0.001). To gain more insight into a gain-of-function hypothesis, we modelled the 3D structure of the full-length protein sequences, a trypsin-like serine protease composed of two folded beta barrels connected by a long loop positioned at the back of the active cleft^63^, and, by adopting a new graph-based theory approach, we inferred the key residues driving the structural differences among loci. Notably, the methodology clustered all the structures into four groups, consistent with the phylogenetic analysis, plus a fifth group for the Melpomene/Silvaniform *Coc2* sub-clade, which evolved under diversifying positive selection. We found that seven residues drive the overall clustering, and these lie in three regions of the 3D structures, corresponding to three loops in regions highly exposed to the solvent (Fig. 7d). The structural alignment of the predicted cocoonases with the X-Ray structures of several homologous human serine proteases (Supplementary Fig. 49), shows that the two largest loops (pos: 217-217 and pos: 119-122) corresponds to highly flexible regions in the experimental structures (*i*.*e*., B-factor), in contrast with the shorter third loop (pos: 68-71), which is in turn analogous to a region with higher stability. These analyses suggest that the duplicated genes might have gained the capacity to bind and process different substrates by changing their flexibility throughout the radiation of Heliconiini. This is consistent with the hypothesis that, in order to obtain a gain-of-function and to give rise to new interactions, a protein needs to change few sites in intrinsically disordered regions^65^. Our combined results present a more complex story than previously described, and both the high copy number variation and patterns of selection within Heliconiinae appear inconsistent with these genes playing a critical role in the evolution of pollen feeding.

**Fig. 7.**
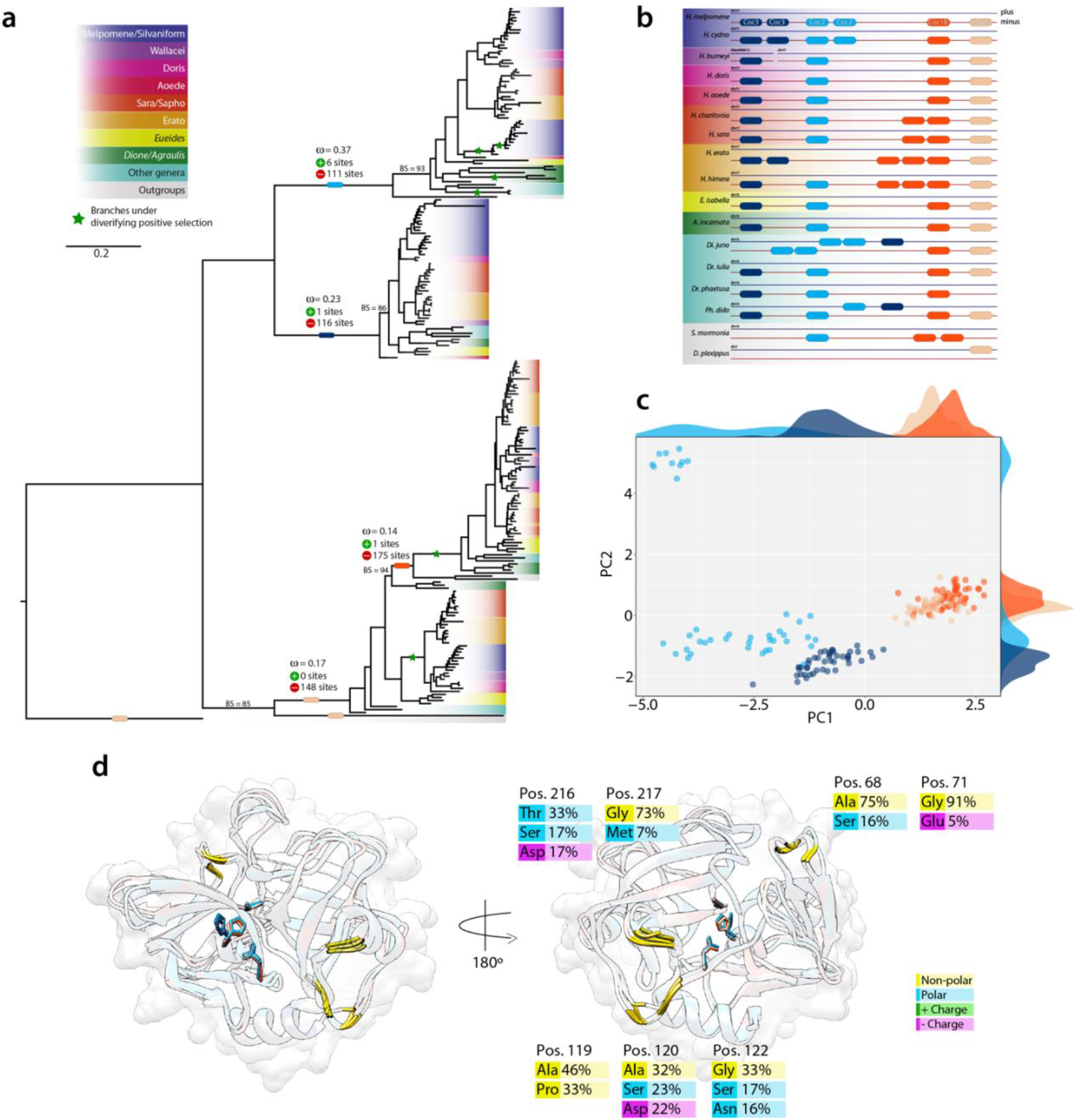
Cocoonase evolution and structural divergence across Heliconiini. **a** Maximum likelihood phylogeny of nucleotide sequences of the four cocoonase loci across Heliconiinae. Coloured blocks on branches show the stem for each locus. The green star indicates branch under diversifying positive selection. Bootstrap (BS) values for main branches are listed for those with values below 95 **b** Synteny map of the different loci for the genomes with highest contiguity. **c** PCA of the network-based analysis of 181 predicted protein models. **d** Structural alignment of the closest sequence to the centroid of each of the five clusters of the PCA (coloured ribbons). For each of structure the three active sites of the active cleft are depicted as sticks, while in gold the seven identified positions the best explains the clustering of the PCA. On the left the same structure rotated 180º. For each of the positions the most frequent amino acids are shown with their respective frequency in the alignment.

## Conclusions

We have curated available genomic data and new reference genomes to build a tribe-wide dataset for Heliconiini butterflies. Using the resulting phylogenetic framework, we examined patterns of genomic change at points in the species tree around which key phenotypic innovations are expected. We investigated the evolution of genome size, its effect on protein-coding gene expansions and contraction, and selective forces such as diversifying positive selection on protein coding genes and the acceleration of conserved non-coding genes. Supported by the characterization of all these genomic features, our analyses ultimately allowed us to unpick the molecular architecture of key innovations in this enigmatic group of butterflies. This provides a genome-wide perspective of the strong but gradual selection events that occurred at the basal branches of the Heliconiini tribe, exemplified by expansions in gene families and OGs linked to biochemical processes relevant to cyanogenic defences, dietary shifts, and longevity, with signatures of adaptive evolution. Notably, multiple strands of evidence implicate selection acting on both coding and non-coding loci affecting neural development and proliferation, synaptic processes, and long-term memory, in line with evidence of substantial variation in the structure of Heliconiini brains^10,11,25,54^. These results highlight how individual loci, as well as wider pathways, such as the Notch and Hippo pathways, might have evolved under a strong diversifying selection, providing the first gene-phenotype links underpinning mushroom body expansion^25^. Finally, our test for acceleration of putative *cis*-regulatory elements (CNEEs) at the stem of *Heliconius*, used for the first time in insects outside the *Drosophila* system, identified more prevalent positive selection on non-coding elements compared to protein coding genes at the origin of *Heliconius*. This suggests the suite of derived phenotypes in this genus might have largely evolved through changes in gene expression via modification of regulatory elements (*e*.*g*.: promoters, enhancers, and silencers)^40,66^. In conclusion, our work offers a comprehensive view to the evolutionary history of an enigmatic tribe of butterflies, the evolution of their genomic architectures, and provides the most thorough analysis of potential molecular changes linked to the physiological and behavioural innovations of a diverse group of butterflies. These gene-phenotype hypothesis, alongside our comprehensive dataset, provide new opportunities to test and derive causative links between molecular and trait innovations.

## Methods Summary

### DNA and RNA Extraction and Sequencing

Individuals of *Dryadula phaetusa, Dione juno, Agraulis vanilla vanillae*, were collected from partially inbred commercial stocks (Costa Rica Entomological Supplies, Alajuela, Costa Rica); individuals of *Agraulis vanilla incarnata* collected from Shady oak butterfly farm (Brooker, Florida, USA); while individuals of *Speyeria mormonia washingtonia* (Washington, USA), *Philaethria dido* (Gamboa, Panama), *Podothricha telesiphe* (Cocachimba, Peru), *H. aoede* (Tarapoto, Peru), *H. doris* (Gamboa, Panama), and *H. cydno*, were collected from the wild. Samples collected in Peru were obtained under permits 0289-2014-MINAGRI-DGFFS/DGEFFS, 020-014/GRSM/PEHCBM/DMA/ACR-CE, 040– 2015/GRSM/PEHCBM/DMA/ACR-CE, granted to Dr Neil Rosser, and samples from Panama were collected under permits SEX/A-3-12, SE/A-7-13 and SE/AP-14-18. High-molecular-weight genomic DNA was extracted from pupae (commercial stock specimens) and adults (wild caught specimens), dissecting up to 100 mg of tissue, snap frozen in liquid nitrogen and homogenized in 9.2 ml buffer G2 (Qiagen Midi Prep Kit). The samples were then transferred to a 15 ml tube and processed with a Qiagen Midi Prep Kit (Qiagen, Valencia, CA) following the manufacturer’s instructions. From the same stocks, RNA was extracted separately from six adult and early ommochrome stage pupae. Each tissue was frozen in liquid nitrogen and quickly homogenized in 500 μl Trizol, adding the remaining 500 μl Trizol at the end of the homogenization. Phase separation was performed by adding 200 μl of cold chloroform. The upper phase was then transferred to RNeasy Mini spin column and processed with a Qiagen RNeasy Mini Prep Kit (Qiagen, Valencia, CA), before DNAse purification using the Turbo DNA-free kit (Life Technologies, Carlsbad, CA) following the manufacturer’s instructions. According to HWM genomic DNA quality, samples were sequenced with PacBio Circular Long Reads, PacBioHiFi, Oxford Nanopore Technology reads, or 10X Genomics Linked Reads, adding librarie s of Illumina DNAseq (HiSeq2500 150 × 2) data for PacBio Circular Long Reads libraries for error corrections. Short-read data for 39 other species where also downloaded from NCBI and used for *de novo* assembly and/or to curate already available assemblies (for further details see Supplementary Methods).

### Long Read *De Novo* Genome Assembly

PacBio Hifi CCS reads were assembled using hifiasm v0.12-r304^67^; while regular PacBio Circular Long Reads were corrected, trimmed and assembled using CANU v1.8 + 356^68^ as in Cicconardi et al. (2021). Resulting assemblies were subsequently corrected with their uncorrected raw PacBio long reads using pbmm2 v1.0.0 and Arrow v2.3.3. Further error correct was performed with short Illumina reads using Pilon v1.23^69^ with five iterations. Mis-assemblies were corrected with Polar Star. Sequenced Illumina paired-end reads from 10X Genomics libraries were input to the Supernova v2.1.1 assembler (10x Genomics, San Francisco, CA, USA)^70^ for *de novo* genome assembly, following optimisation of parameters. Tigmint v1.1.2^71^ with default settings was then adopted to identify misassemblies. The final step of scaffolding was performed using ARCS v1.1.0^72^, a scaffolding procedure that utilizes the barcoding information contained in linked-reads to further organize assemblies. In all assemblies described thus far, haplocontigs were removed using Purge Haplotigs v20191008^73^, and PacBio and Nanopore data (when available) were used to perform the first stage of scaffolding with LRScaf v1.1.5^74^. If RNA-seq data were available, P_RNA_scaffolder was also used to further scaffolding. Gap filling was performed with LR_Gapcloser v.1.1^75^. Before the chromosome-level scaffolding, we used synteny maps implemented with BLAST ^76^ and ALLMAPS^77^ to identify duplicated regions at the end of scaffolds, manually curating the scaffolds to trim them away. Duplication level and completeness were checked with BUSCO (Benchmarking Universal Single-Copy Orthologs; v3.1.0, Insecta_odb9)^78^ at each step of the assembling to keep track of losses and fragmentation of genomic regions.

### Reference-Based Genome Assembly

To assemble the genomes from the retrieved Illumina PE reads from NCBI (see Supplementary Table 1) we implemented a reference-guided assembly approach, adapting and extending the protocol from Lischer and Shimizu (2017). The strategy involves first mapping reads against a reference genome of a related species (see Supplementary Table 1 ‘*Ref Genome*’ field) to reduce the complexity of the *de novo* assembly within continuous covered regions, then integrating reads with no similarity to the related genome in a further step. Modifications from the reference guided *de novo* pipeline^79^ started in the 1^st^ step, were paired-end Illumina reads were mapped onto the reference genome with Minimap2 v2.17-r974-dirty^80^, followed by Pilon^69^. This procedure increases the mapability of the target species to the reference by modifying the reference assembly at the nucleotide level. Paired-end reads were then mapped against the modified version of the reference assembly Bowtie2 v2.2.1^81^, and assigned into blocks as in the original pipeline. For each block, reads were *de novo* assembled using SPAdes v3.15^82^. Redundancy generated at this stage was removed as in the original pipeline. After the final step, short-reads were used to attempt scaffolding and gap closing with SOAPdenovo2 vr240^83^. Leveraging the very small genetic differences between these species and their reference genomes, a final assembly scaffolding was performed with RaGOO^84^, a homology-based scaffolding and misassembly correction pipeline. RaGOO^84^ identifies structural variants and sequencing gaps, to accurately orders and orient *de novo* genome assemblies. Abyss-sealer v2.2.2 from the Abyss package^85^ was use as last step to attempt to close remaining gaps.

### Curation of Available Illumina Assemblies

Available Heliconiini assemblies from Edelman et al. (2019) were included in our dataset with a small, but effective curation. We checked for contaminants, as for the previous *de novo* and reference guided assemblies (see below), and at the haplocontig level (using BUSCO, see above). Raw Illumina reads were remapped onto their own assembly and Purge Haplotigs, with ad hoc -a parameter, was adopted to remove haplocontigs, followed by a scaffolding procedure with SOAPdenovo2 (127mer). A synteny map was generated with ALLMAPS and, using their closest available reference assembly, a chromosomal scaffolding was generated. This procedure was adopted to maximise the contiguity. This represents a similar procedure to that recently adopted to scaffolded draft genomes^87^. Finally, Abyss-sealer v2.2.2 from the Abyss package^85^ was used for gap filling (see above).

### Bacterial Contamination & Assembly Completeness Assessment

After the genome assembly stage all datasets were analysed to remove contaminants. Blobtools v1.1.1^88^ was used to filter out any scaffolds and contigs assigned to fungal and bacterial contaminants. Furthermore, mitochondrial sequences were identified by blasting (BLASTn) contigs and scaffolds against the mitochondrial genome of available *Heliconius ssp*.. Finally a combination of BUSCO^78^, with the Insecta set in OrthoDB v.9 [-m genome], and Exonerate v2.46.2^89^, was used to assess genome completeness and duplicated content.

### Whole Genome Alignment & Genome Evolution

BUSCO complete single-copy orthologous genes were used to generate a first draft of the phylogeny to guide the whole genome alignment. The nucleotide sequence of each locus aligned with MACSE v2.03^90^ and concatenated into a single alignment. A maximum-likelihood (ML) search was adopted to estimate the phylogenetic tree as implemented in FastTree v2.1.11 SSE3^91^. All 63 soft-masked genomes were then aligned with Cactus v1.2.3^92,93^ with chromosome-level genomes as reference. Post-processing was performed by extracting information from the resulting hierarchical alignment (HAL). As a measure of genome size, we adopted the assembly size. Although this approach has some limitations, the high BUSCO scores, and the lack of correlation between assembly size and contiguity (R^2^ = 0.002; ρ = 0.05) indicates that the great majority of the assemblies are complete, most of the smaller assembly sizes are unlikely to be artifacts of incomplete assembly, and the quality control during assembly ensured that larger genomes were not due to DNA contamination. Therefore, assembly size should closely correlate with the actual genome size, and no circularity or biases should be present. We then used HALSummarizeMutations, from the Cactus package, to summarize inferred mutations at each branch of the underlying Nymphalid phylogeny. We calculated rates for transposition (d_*P*_), insertion (d_*I*_), deletion (d_*D*_), inversion (d_*V*_), and duplication (d_*U*_) events per million years (Ma) of evolution, based on the inferred divergence estimates from the phylogeny (see below). Ancestral state reconstruction of genome size was assessed using the maximum likelihood method implemented in the R package phytools^94^. Evolutionary conservation at individual alignment sites, phyloP scores (CONACC) were computed using a neutral model as implemented in halPhyloPTrain.py script (Cactus package). A non-overlapped sliding window of 10bp was adopted and data partitioned according to coding, intronic, 5’UTR and 3’UTR regions (further details see Supplementary Materials & Methods).

### Gene Prediction and Transcriptome Annotation

The NCBI SRA archive was explored, and the best SRA archives were downloaded, based on abundance and tissue (Supplementary Table 2). These were used to re-annotate genes for their reference genomes (*e*.*g*., *H. erato v. 1* and *H. melpomene v*.*2*.*5*). Short-reads were quality filtered and trimmed Trimmomatic V0.39^95^, and predicted coding genes, *ab initio* and *de novo* approaches were implemented and combined in a pipeline with the aim of obtaining the best from each approach to overcome their own limitations. Reads from multiple tissues (when available) were pooled, mapped with STAR v2.7.10a^96^ and used as training data for the BRAKER v2.1.5 pipeline^97^, using AUGUSTUS v3.4.0^98^, along with the masked genomes generated with RepeatMasker v4.1.1^99^, using the Lepidoptera database. The gene predictions were followed by the UTR annotation step via GUSHR v1.1.0 (Gaius-Augustus/GUSHR, 2020). The *de novo* transcriptome assemblies were generated using Trinity v2.10.0^100,101^ separately for each tissue. To generate the *ab initio* transcriptomes, tissue-specific reads were realigned to the genome using STAR and assembled using both Stringtie v2.1.3b^102^ and Cufflinks v2.2.1^103–105^. The BAM files were used as input for Portcullis v1.1.2^106^ to validate splice-site DBs and together with the previously generate four types of annotations (prediction, *de novo*, two *ab initio*) were combined using Mikado v2.3.3^107^. Finally, we used the Comparative Annotation Toolkit (CAT)^108^, a comparative annotation pipeline that combines a variety of parameterizations of AUGUSTUS, including Comparative AUGUSTUS, with transMap projections, to annotate all the species through the whole-genome Cactus alignments to produce an annotation set on every genome in alignment using *E. isabella* as a reference species (further details see Supplementary Materials & Methods).

### Intron Evolution Analyses

Intronic regions were extracted from the longest transcript of each gene model using the annotations, as in Cicconardi et al^19^. Sequences were scanned with RepeatMasker. For each species introns were divided into short and long based on their median values. Their relative scaling coefficients and intercepts were subsequently analysed with SMATR^109^. The intron turnover rate was subsequently estimated using Malin (Mac OS X version)^110^ to infer their conservation status in ancestral nodes, and the turnover rate (gain/loss) at each node with Malin’S built-in model ML optimization procedure (1,000 bootstrap iterations) (further details see Supplementary Materials & Methods).

### Functional Annotation and Orthologous-Group Dynamic Evolution

Orthology inference was performed as implemented in Broccoli v1.1^111^ optimizing parameters (Romain Derelle, personal communication) for a more reliable list of single-copy orthologous groups (scOGs). Broccoli also returns a list of chimeric transcripts, which were manually curated in the *E. isabella* transcriptome, with a set of custom scripts that were implemented to automate the process in all the other taxa (BroccoliChimeraSplitDataGather.py; available at https://francicco@bitbucket.org/ebablab/custum-scripts.git), before a second Broccoli iteration. From each OG, a putative functional annotation was performed by identifying both the protein domain architecture using HMMER v3.3.2 (HMMScan)^112^ with DAMA v2^113^. Annotation of GO terms were assigned with a homology-based search against *Drosophila melanogaster* protein databases (FlyBase.org), and with a predictive-based strategy with CATH assignments^114–116^, scanning against the library of CATH functional family (FUNFAMS v4.2.0) HMMs^116^. We modelled OG expansions and contractions as implemented in CAFE v5.0 using only genomes with complete BUSCO genes ≥ 90% (52/64 species) (further details see Supplementary Materials & Methods).

### Phylogenetic Analysis & Divergence Estimates

Fully processed alignments of scOG were selected, concatenated and used to generate a maximum likelihood (ML) phylogenetic tree, as implemented in IQ-Tree2, partitioning the supermatrix for each locus and codon position, and with 5,000 ultrafast bootstrap replicates, resampling partitions, and then sites within resampled partitions^117,118^. ILS was explored performing a coalescent summary method species tree using scOG gene trees, as implemented in ASTRAL-III v5.6.3^119^. To further explore phylogenetic support, the quartet sampling (QS) analysis was performed^120^. The Bayesian algorithm of MCMCtree v4.8a (from PAML package)^121^ with approximate likelihood computation was used to estimate divergence times for the whole dataset. Branch lengths were estimated by ML and then the gradient and Hessian matrix around these ML estimates were computed under MCMCtree using the DNA supermatrix. From the TimeTree database^122^, four calibration points with uniform distributions were used (supplementary table 3), consistent with previous phylogenetic studies of Heliconiini^8,122^. For these priors a birth-death process with λ=μ=1 and ρ=0, and a diffuse gamma-Dirichlet priors was given for the molecular rate (Γ=2,20) and a diffusion rate (σ2=2,2). Ten independent runs were executed, each with a burn-in of 2,500,000 generations. Convergence was checked using Tracer V1.7.1^123^ (further details see Supplementary Materials & Methods).

### Genome-Wide Scan for Introgression

Patterns of introgression within Heliconiini were scanned using discordant-count test (DCT) and the branch-length test (BLT), which rely on the topologies of gene trees for triplets of species as implemented in Suvorov et al.^14^. These tests were applied on all triplets extracted from scOG gene trees within Heliconiini, and the resulting *P*-values were then corrected for multiple testing using the Benjamini-Hochberg procedure with a false discovery rate (FDR) cut-off of 0.05. Dsuite^124^ was then used to plot the results in a heatmap plot^14^ (further details see Supplementary Materials & Methods).

### Evolution of Gene Families

Fifty-seven gene families spanning receptors, enzymes, channels, and transporters were selected to further explore the evolution of Heliconiini (Supplementary Table 4). Amino acid sequences of the entire gene family (GF) were aligned using CLUSTALW v1.2.1^125^ and used to build a ML tree using FastTree v 2.1.11 SSE3 and used as input for MIPhy v1.1.2^126^, in order to automatically predict members of orthologous groups for each GFs, leveraging a species tree. For each GF, OG copy number was processed with CAFE (see above) to further explore events of expansion and contraction. From each OG in-paralogs were removed (custom python script RemoveInParalogFromTree.py available at https://francicco@bitbucket.org/ebablab/custum-scripts.git). If the procedure generated a single-copy OG (nscOGs) it was analysed by contrasting evolutionary pressures between *Eueides* and *Heliconius* species. The signature of selection (aBSREL) and relaxation (RELAX^127^) were performed as implemented in HyPhy (further details see Supplementary Materials & Methods).

### Diversifying Positive Selection Across Heliconiini

Evolutionary trajectories across Heliconiini were performed with a pipeline similar to that in Cicconardi *et al*.^3,128,129^ computing *ω* (the ratio of nonsynonymous to synonymous substitution rates; d_*N*_/d_*S*_) on five branches of the Nymphalid phylogeny using codon-based alignments of groups of one-to-one orthologs (scOGs). Compared to previous pipelines, we introduced a pre-alignment filtering procedure as implemented in PREQUAL v1.02^130^, and a post-alignment filtering with HmmCleaner^131^. A ML gene tree was then generated as implemented in IQ-Tree2 v2.1.3 COVID-edition and the adaptive branch-site random effects likelihood (aBSREL) method^132,133^ used, as implemented in the HyPhy batch language^134^. Enrichment of GOterms was performed using a combination of two different approaches, the hyperGTest algorithm, implemented in the GOStats package^135^ for R and GOATOOLS^136^, and only considering significant terms in common between GOStats and GOATOOLS were considered (*P*-value < 0.05). The GeneMANIA prediction server^137–139^ was used to predict functions of genes under selection (FDR cut-off of 0.05) (further details see Supplementary Materials & Methods).

### Accelerated CNEEs

Conserved non-exonic elements (CNEEs) where annotated from the 63-way whole genome alignment using the phast V1.4 package^140,141^, using the *E. isabella* as reference. Elements from the first round of annotation were merged if they were within 5 bp of each other into single conserved element, and regions shorter than 50 bp, with less than 50 species, and gaps in more than 50% of the consensus, excluded. Acceleration on the *Heliconius* stem was tested in a Bayesian framework as implemented in PhyloAcc-GT^42^. We considered CNEEs that had the BF1 >= 10, and BF2 >= 1, specifically on the *Heliconius* stem. Gene-wise and Special-wise enrichment were computed with 10,000 randomly resamples of the entire list of CNEEs and tested with a binomial test (*Pr*_*binomial*_ = observed aCNEEs per gene/region | number of aCNEEs, expected aCNEEs per gene/region]). To test for gene ontology terms (GO) of functional elements enriched in *Heliconius*-accelerated CNEEs, we used two permutation approaches and a genomic fraction approach. All account for possible biases towards particular gene functions or CNEE distributions (further details see Supplementary Materials & Methods).

### Cocoonase Annotation & Analysis

The protein sequences of *Heliconius* cocoonases were obtained from Smith et al.^63^, while the sequence from *Bombyx mori* was downloaded from NCBI. Sequences were used as queries for map protein sequences onto all the 63 assemblies, using Exonerate, and subsequently manually corrected. All nucleotide sequences were quality filtered using Prequal aligned using MACSE, to generate a single ML gene tree adopting IQ-Tree2. All sequences from each of the four clades were realigned separately and several tests were implemented in HyPhy (overall *ω*; SLAC; aBSREL; RELAX). In particular, the sign of diversifying positive selection (aBSREL) was detected by scanning all internal branches of the whole cocoonases phylogeny, correcting for multiple tests using a final P-value threshold of 0.05. The structural analyses were conducted on the 181 full length sequences (∼ 220 aa in length), removing peptide signal detected with SignalP v5.0b^142^, and aligning sequences with the closest homolog protein for which a crystal structure is available (pdb: 4AG1), identified by HHpred server^143^. To predict 3D structures the RoseTTAFold v1.1.0 pipeline^144^ was adopted. A graph theory based analysis was performed for each 3D model belonging to the final data set, as implemented in Ruiz-Serra et al.^145^. The method adopts graph-based metrics to capture both the local features of the predicted distance maps (strength) as well as to characterize global patterns of the molecular interaction network. After performing a multiple alignment among all the sequences of the data set, we obtained an *N* x *M* output matrix, where *N* is the total number of the sequences and *M* the total number of all residue position, and used to performed a Principal Component Analysis (PCA) with the aim of projecting each *M*-dimensional vector (i.e., the set of strength values associated to each protein of the data set) into essential space (i.e., the PCA space). Consistency between the phylogenetic signal and the structural information of the four loci was evaluated by checking how groups are separated from each other in the PCA space, calculating four distributions of both the first and second component, and performing a Kolmogorov-Smirnov (K-S) test^146^ as implemented in the R function ks.test. Key residues were identified selecting the residues that explained the most fraction of the first two PCA components(further details see Supplementary Materials & Methods).

## Data Availability

Data and code used for these analyses are available on NCBI under their project id (see Supplementary table 1) and GitHub (https://github.com/francicco/-ComparativeGenomicsOfHeliconiini). Note: for submission reasons it was not possible to attach the Supplementary tables, these can be download on the GitHub repository (natcom_supplementarytables.xlsx.zip) Individual mark-recapture datasets can be obtained by contacting specific dataset owners.

## ACKNOWLEDGMENTS

This article would not be possible without the great support of the great *Heliconius* community. We are also grateful to the environmental ministries in Peru and Panama for permission to collect and export samples and the STRI community for assistance in the field. F.C. would like to thank Ronald Mori Pezo for his great help in collecting *H. aoede* and *Podotricha telesiphe*; Angel Corpuz for various informatics support including a great patience; Ian Fiddes, Mark Diekhans, Glen Hichey and Marina Haukness for their great support for Cactus and CAT; Gregg Thomas and Tim Sackton for they help with PhyloACC-ST, preparation and analysis; Federica Cattonaro, Davide Scaglione and Simone Scalabrin from IGA (Udine, Italy) for their fruitful discussion on the best sequencing strategy to perfom. F.C. and S.H.M. are grateful to the High-Performance Computing team at the Advanced Computing Research Centre, University of Bristol for support. **Funding:** This article was supported by NERC IRF (NE/N014936/1) and ERC Starter Grant (758508) to S.H.M. S.M. was supported by the Royal Society URF\R1\180682. F.C. was supported as a postdoctoral researcher ERC.

## Author contributions

Conceptualization: S.H.M and F.C. Data collection: All authors. Genomic analysis: F.C. Structural analyses: F.C., E.M. and D.d.M. Visualization: F.C. Funding acquisition: S.H.M. Writing – original draft: F.C. and S.H.M. Writing – review & editing: All authors.

## Competing interests

The authors declare that they have no competing interests.

**Correspondence** and requests for materials should be addressed to Francesco Cicconardi and Stephen Montgomery.

## References

1. Stroud, J. T. & Losos, J. B. Ecological Opportunity and Adaptive Radiation. Annu. Rev. Ecol. Evol. Syst. 47, 507–532 (2016).

2. Erwin, D. H. A conceptual framework of evolutionary novelty and innovation. Biol. Rev. 96, 1–15 (2021).

3. Cicconardi, F. et al. Genomic signature of shifts in selection in a subalpine ant and its physiological adaptations. Mol. Biol. Evol. 1–17 (2020) doi:10.1093/molbev/msaa076.

4. Yuan, Y. et al. Comparative genomics provides insights into the aquatic adaptations of mammals. Proc. Natl. Acad. Sci. U. S. A. 118, 1–9 (2021).

5. Kozak, K. M., Joron, M., McMillan, W. O. & Jiggins, C. D. Rampant Genome-Wide Admixture across the Heliconius Radiation. Genome Biol. Evol. 13, 1–17 (2021).

6. Martin, S. H., Davey, J. W., Salazar, C. & Jiggins, C. D. Recombination rate variation shapes barriers to introgression across butterfly genomes. PLoS Biol. (2019) doi:10.1371/journal.pbio.2006288.

7. Edelman, N. B. et al. Genomic architecture and introgression shape a butterfly radiation. 599, 594–599 (2019).

8. Kozak, K. M. et al. Multilocus species trees show the recent adaptive radiation of the mimetic heliconius butterflies. Syst. Biol. 64, 505–524 (2015).

9. de Castro, É. C. P. et al. Sequestration and biosynthesis of cyanogenic glucosides in passion vine butterflies and consequences for the diversification of their host plants. Ecol. Evol. 9, 5079–5093 (2019).

10. Young, F. J. & Montgomery, S. H. Pollen feeding in Heliconius butterflies : the singular evolution of an adaptive suite. Proc. R. Soc. B Biol. Sci. 287, (2020).

11. Montgomery, S. H., Merrill, R. M. & Ott, S. R. Brain composition in Heliconius butterflies, posteclosion growth and experience-dependent neuropil plasticity. J. Comp. Neurol. 524, 1747–1769 (2016).

12. Hawornwattana, Y. U. T., Eixas, F. E. A. S., Ang, Z. I. Y. & Allet, J. A. M. Full-Likelihood Genomic Analysis Clarifies a Complex History of Species Divergence and Introgression : The Example of the erato − sara Group of Heliconius Butterflies. 71, 1159–1177 (2022).

13. Thawornwattana, Y., Seixas, F. A., Yang, Z. & Mallet, J. Full-Likelihood Genomic Analysis Clarifies a Complex History of Species Divergence and Introgression: The Example of the erato-sara Group of Heliconius Butterflies. Syst. Biol. 71, 1159–1177 (2022).

14. Suvorov, A. et al. Widespread introgression across a phylogeny of 155 Drosophila genomes. Curr. Biol. 1–13 (2021) doi:10.1016/j.cub.2021.10.052.

15. Walters, J. R., Corbins, C., Hardcastle, T. J. & Jiggins, C. D. Evaluating female remating rates in light of spermatophore degradation in Heliconius butterflies: Pupal-mating monandry versus adult-mating polyandry. Ecol. Entomol. 37, 257–268 (2012).

16. Thurman, T. J., Brodie, E., Evans, E. & McMillan, W. O. Facultative pupal mating in Heliconius erato: Implications for mate choice, female preference, and speciation. Ecol. Evol. 8, 1882–1889 (2018).

17. Kapusta, A., Suh, A. & Feschotte, C. Dynamics of genome size evolution in birds and mammals. Proc. Natl. Acad. Sci. U. S. A. (2017) doi:10.1073/pnas.1616702114.

18. Ruggieri, A. A. et al. A butterfly pan-genome reveals a large amount of structural variation underlies the evolution of chromatin accessibility. bioRxiv 2022.04.14.488334 (2022).

19. Cicconardi, F. et al. Chromosome Fusion Affects Genetic Diversity and Evolutionary Turnover of Functional Loci but Consistently Depends on Chromosome Size. Mol. Biol. Evol. 38, 4449–4462 (2021).

20. Sun, C. et al. Genus-Wide Characterization of Bumblebee Genomes Provides Insights into Their Evolution and Variation in Ecological and Behavioral Traits. Mol. Biol. Evol. 38, 486–501 (2021).

21. Neafsey, D. E. et al. Highly evolvable malaria vectors: The genomes of 16 Anopheles mosquitoes. Science (80-.). 347, (2015).

22. de Castro, É. C. P., Musgrove, J., Bak, S., McMillan, W. O. & Jiggins, C. D. Phenotypic plasticity in chemical defence allows butterflies to diversify host use strategies. bioRxiv (2020) doi:10.1101/2020.04.07.030122.

23. Pinheiro de Castro, É. C. et al. The dynamics of cyanide defences in the life cycle of an aposematic butterfly: Biosynthesis versus sequestration. Insect Biochem. Mol. Biol. 116, 103259 (2020).

24. Du, M. et al. Identification of lipases involved in PBAN stimulated Pheromone production in Bombyx mori using the DGE and RNAi approaches. PLoS One 7, (2012).

25. Couto, A. et al. Rapid expansion and visual specialization of learning and memory centers in Heliconiini butterflies. bioRxiv (2022).

26. Mendes, F. K., Vanderpool, D., Fulton, B. & Hahn, M. W. CAFE 5 models variation in evolutionary rates among gene families. Bioinformatics 36, 5516–5518 (2020).

27. Lage, J. L. Da, Thomas, G. W. C., Bonneau, M. & Courtier-Orgogozo, V. Evolution of salivary glue genes in Drosophila species. BMC Evol. Biol. 9, (2018).

28. Opitz, S. E. W. & Müller, C. Plant chemistry and insect sequestration. Chemoecology 19, 117–154 (2009).

29. Sung, E. J. et al. Cytokine signaling through Drosophila Mthl10 ties lifespan to environmental stress. Proc. Natl. Acad. Sci. U. S. A. (2017) doi:10.1073/pnas.1712453115.

30. Wu, L. et al. CYP303A1 has a conserved function in adult eclosion in Locusta migratoria and Drosophila melanogaster. Insect Biochem. Mol. Biol. 113, 103210 (2019).

31. Tang, B., Wang, S. & Zhang, F. Two storage hexamerins from the beet armyworm Spodoptera exigua: Cloning, characterization and the effect of gene silencing on survival. BMC Mol. Biol. 11, (2010).

32. Portin, P. & Portin, P. General outlines of the molecular genetics of the Notch signalling pathway in Drosophila melanogaster: A review. Hereditas 136, 89–96 (2002).

33. Li, X., Xie, Y. & Zhu, S. Notch maintains Drosophila type II neuroblasts by suppressing expression of the fez transcription factor earmuff. Dev. 143, 2511–2521 (2016).

34. Sackton, T. B. et al. Convergent regulatory evolution and the origin of flightlessness in palaeognathous birds. Science (80-.). 364, 74–78 (2019).

35. Lin, Q. et al. The seahorse genome and the evolution of its specialized morphology. Nature 540, 395–399 (2016).

36. Snetkova, V., Pennacchio, L. A., Visel, A. & Dickel, D. E. Perfect and imperfect views of ultraconserved sequences. Nat. Rev. Genet. 23, 182–194 (2022).

37. McLean, C. Y. et al. Human-specific loss of regulatory DNA and the evolution of human-specific traits. Nature 471, 216–219 (2011).

38. Sackton, T. B. et al. Convergent regulatory evolution and loss of flight in paleognathous birds. Science (80-.). 364, 74–78 (2019).

39. Pollard, K. S., Hubisz, M. J., Rosenbloom, K. R. & Siepel, A. Detection of nonneutral substitution rates on mammalian phylogenies. Genome Res. 20, 110–121 (2010).

40. Van Belleghem, S. M. et al. High level of novelty under the hood of convergent evolution. Science 379, 1043–1049 (2023).

41. Hu, Z., Sackton, T. B., Edwards, S. V. & Liu, J. S. Bayesian Detection of Convergent Rate Changes of Conserved Noncoding Elements on Phylogenetic Trees. Mol. Biol. Evol. 36, 1086–1100 (2019).

42. Han Yan et al. PhyloAcc-GT: A Bayesian method for inferring patterns of substitution rate shifts and associations with binary traits under gene tree discordance. (2022).

43. Roblodowski, C. & He, Q. Drosophila Dunc-115 mediates axon projection through actin binding. Invertebr. Neurosci. 17, (2017).

44. Frank, C. A. & James, T. D. Homeostatic control of Drosophila neuromuscular junction function. 1–13 (2020) doi:10.1002/syn.22133.

45. Heymann, C. et al. Molecular insights into the axon guidance molecules Sidestep and Beaten path. 1–18 (2022) doi:10.3389/fphys.2022.1057413.

46. Chen, K., Richlitzki, A., Featherstone, D. E., Schwärzel, M. & Richmond, J. E. Tomosyn-dependent regulation of synaptic transmission is required for a late phase of associative odor memory. (2011) doi:10.1073/pnas.1110184108.

47. Protection, N., Drosophila, A., Hospital, W., Shcool, H. M. & Hughes, H. An Evolutionarily Conserved Role of Presenilin in. 206, 1479–1493 (2017).

48. Sun, J., Zhang, J., Wang, D. & Shen, J. The transcription factor Spalt and human homologue SALL4 induce cell invasion via the dMyc-JNK pathway in Drosophila. 1, 1–10 (2020).

49. Closser, M. et al. Article An expansion of the non-coding genome and its regulatory potential underlies vertebrate neuronal diversity ll Article An expansion of the non-coding genome and its regulatory potential underlies vertebrate neuronal diversity. Neuron 110, 70-85.e6 (2022).

50. Link, B. A. The Roles of Hippo Signaling Transducers Yap and Taz in Chromatin Remodeling. (2019).

51. Stem, G. & Progeny, C. The Osa-Containing SWI/SNF Chromatin-Remodeling Complex Is Required in the Germline Differentiation Niche for Germline Stem Cell Progeny Differentiation. 1–19 (2021).

52. Chubak, M. C. et al. Individual components of the SWI / SNF chromatin remodelling complex have distinct roles in memory neurons of the Drosophila mushroom body. (2019) doi:10.1242/dmm.037325.

53. Farris, S. M. Evolution of complex higher brain centers and behaviors: Behavioral correlates of mushroom body elaboration in insects. Brain. Behav. Evol. 82, 9–18 (2013).

54. Couto, A., Young, F. & Stephen, M. Muschroom body expantion in Heliconiini. prep.

55. Sahu, M. R. & Mondal, A. C. Neuronal Hippo signaling: From development to diseases. Dev. Neurobiol. 81, 92–109 (2021).

56. Kaya-çopur, A. et al. The hippo pathway controls myofibril assembly and muscle fiber growth by regulating sarcomeric gene expression. Elife 10, 1–34 (2021).

57. Abeysundara, N., Simmonds, A. J. & Hughes, S. C. Moesin is involved in polarity maintenance and cortical remodeling during asymmetric cell division. Mol. Biol. Cell 29, 419–434 (2018).

58. Wang, X., Zhang, Y. & Blair, S. S. Fat-regulated adaptor protein Dlish binds the growth suppressor Expanded and controls its stability and ubiquitination. Proc. Natl. Acad. Sci. U. S. A. 116, 1319–1324 (2019).

59. Bahrampour, S. & Thor, S. Ctr9, a key component of the paf1 complex, affects proliferation and terminal differentiation in the developing drosophila nervous system. G3 Genes, Genomes, Genet. 6, 3229–3239 (2016).

60. Loyer, N. & Januschke, J. Where does asymmetry come from? Illustrating principles of polarity and asymmetry establishment in Drosophila neuroblasts. Curr. Opin. Cell Biol. 62, 70–77 (2020).

61. Blair, S. & McNeill, H. Big roles for Fat cadherins. Curr. Opin. Cell Biol. 51, 73–80 (2018).

62. Yildirim, K., Petri, J., Kottmeier, R. & Klämbt, C. Drosophila glia: Few cell types and many conserved functions. Glia 67, 5–26 (2019).

63. Smith, G. et al. Evolutionary and structural analyses uncover a role for solvent interactions in the diversification of cocoonases in butterflies. Proc. R. Soc. B Biol. Sci. 285, (2018).

64. Gai, T. et al. Cocoonase is indispensable for Lepidoptera insects breaking the sealed cocoon. PLoS Genet. 16, 1–16 (2020).

65. Gerasimavicius, L., Livesey, B. J. & Marsh, J. A. Loss-of-function, gain-of-function and dominant-negative mutations have profoundly different effects on protein structure. 1–15 (2022) doi:10.1038/s41467-022-31686-6.

66. Kaplow, I. M. et al. Relating enhancer genetic variation across mammals to complex phenotypes using machine learning. Science (80-.). 380, 2022.08.26.505436 (2023).

67. Cheng, H., Concepcion, G. T., Feng, X., Zhang, H. & Li, H. Haplotype-resolved de novo assembly using phased assembly graphs with hifiasm. Nat. Methods 18, 170–175 (2021).

68. Koren, S. et al. Canu : scalable and accurate long--read assembly via adaptive k--mer weighting and repeat separation. 1–35 (2016) doi:10.1101/gr.215087.116.Freely.

69. Walker, B. J. et al. Pilon: An integrated tool for comprehensive microbial variant detection and genome assembly improvement. PLoS One 9, (2014).

70. Zheng, G. X. Y. et al. Haplotyping germline and cancer genomes with high-throughput linked-read sequencing. Nat. Biotechnol. 34, 303–311 (2016).

71. Jackman, S. D. et al. Tigmint: Correcting assembly errors using linked reads from large molecules. BMC Bioinformatics 19, 1–10 (2018).

72. Yeo, S., Coombe, L., Warren, R. L., Chu, J. & Birol, I. ARCS: Scaffolding genome drafts with linked reads. Bioinformatics 34, 725–731 (2018).

73. Roach, M. J., Schmidt, S. A. & Borneman, A. R. Purge Haplotigs: Allelic contig reassignment for third-gen diploid genome assemblies. BMC Bioinformatics 19, 1–10 (2018).

74. Qin, M. et al. LRScaf: Improving Draft Genomes Using Long Noisy Reads. bioRxiv 374868 (2018) doi:10.1101/374868.

75. Xu, G. C. et al. LR-Gapcloser: A tiling path-based gap closer that uses long reads to complete genome assembly. Gigascience 8, 1–14 (2018).

76. Camacho, C. et al. BLAST command line applications user manual. (2013).

77. Tang, H. et al. ALLMAPS: Robust scaffold ordering based on multiple maps. Genome Biol. 16, 1–15 (2015).

78. Simão, F. A., Waterhouse, R. M., Ioannidis, P., Kriventseva, E. V. & Zdobnov, E. M. BUSCO: Assessing genome assembly and annotation completeness with single-copy orthologs. Bioinformatics 31, 3210–3212 (2015).

79. Lischer, H. E. L. & Shimizu, K. K. Reference-guided de novo assembly approach improves genome reconstruction for related species. 1–12 (2017) doi:10.1186/s12859-017-1911-6.

80. Li, H. Minimap2: Pairwise alignment for nucleotide sequences. Bioinformatics 34, 3094–3100 (2018).

81. Langmead, B., Trapnell, C., Pop, M. & Salzberg, S. L. Ultrafast and memory-efficient alignment of short DNA sequences to the human genome. Genome Biol. 10, R25 (2009).

82. Bankevich, A. et al. SPAdes: A New Genome Assembly Algorithm and Its Applications to Single-Cell Sequencing. J. Comput. Biol. 19, 455–477 (2012).

83. Luo, R. et al. SOAPdenovo2: an empirically improved memory-efficient short-read de novo assembler. Gigascience 1, 18 (2012).

84. Alonge, M. et al. RaGOO: Fast and accurate reference-guided scaffolding of draft genomes. Genome Biol. 20, 1–17 (2019).

85. Paulino, D. et al. Sealer: A scalable gap-closing application for finishing draft genomes. BMC Bioinformatics 16, 1–8 (2015).

86. Edelman, N. B. et al. Genomic architecture and introgression shape a butterfly radiation. Science (80-.). 366, 594–599 (2019).

87. Seixas, F. A., Edelman, N. B. & Mallet, J. Synteny-Based Genome Assembly for 16 Species of Heliconius Butterflies, and an Assessment of Structural Variation across the Genus. Genome Biol. Evol. 13, 1–18 (2021).

88. Laetsch, D. R. & Blaxter, M. L. BlobTools: Interrogation of genome assemblies. F1000Research 6, 1287 (2017).

89. Slater, G. S. C. & Birney, E. Automated generation of heuristics for biological sequence comparison. BMC Bioinformatics 6, 1–11 (2005).

90. Ranwez, V. et al. MACSE v2: Toolkit for the Alignment of Coding Sequences Accounting for Frameshifts and Stop Codons. Mol. Biol. Evol. 2–4 (2018) doi:10.1093/molbev/msy159.

91. Price, M. N., Dehal, P. S. & Arkin, A. P. FastTree 2 - Approximately maximum-likelihood trees for large alignments. PLoS One 5, e9490 (2010).

92. Armstrong, J. et al. Progressive alignment with Cactus: A multiple-genome aligner for the thousand-genome era. bioRxiv (2019) doi:10.1101/730531.

93. Paten, B. et al. Cactus: Algorithms for genome multiple sequence alignment. Genome Res. 21, 1512–1528 (2011).

94. Revell, L. J. phytools: An R package for phylogenetic comparative biology (and other things). Methods Ecol. Evol. 3, 217–223 (2012).

95. Bolger, A. M., Lohse, M. & Usadel, B. Trimmomatic: A flexible trimmer for Illumina sequence data. Bioinformatics 30, 2114–2120 (2014).

96. Dobin, A. et al. STAR: ultrafast universal RNA-seq aligner. Bioinformatics 29, 15–21 (2013).

97. Brůna, T., Hoff, K. J., Lomsadze, A., Stanke, M. & Borodovsky, M. BRAKER2: automatic eukaryotic genome annotation with GeneMark-EP+ and AUGUSTUS supported by a protein database. NAR Genomics Bioinforma. 3, 1–11 (2021).

98. Stanke, M. et al. AUGUSTUS: A b initio prediction of alternative transcripts. Nucleic Acids Res. 34, 435–439 (2006).

99. Smit, A., Hubley, R. & Green, P. RepeatMasker Open-4.0. 2013–2015. http://www.repeatmasker.org (2013).

100. Iyer, M. K. & Chinnaiyan, A. M. RNA-Seq unleashed. Nat. Biotechnol. 29, 599–600 (2011).

101. Haas, B. J. et al. De novo transcript sequence reconstruction from RNA-seq using the Trinity platform for reference generation and analysis. Nat. Protoc. 8, 1494–512 (2013).

102. Pertea, M. et al. StringTie enables improved reconstruction of a transcriptome from RNA-seq reads. Nat. Biotechnol. 33, 290–295 (2015).

103. Trapnell, C. et al. Differential gene and transcript expression analysis of RNA-seq experiments with TopHat and Cufflinks. Nat. Protoc. 7, 562–578 (2012).

104. Garber, M., Grabherr, M. G., Guttman, M. & Trapnell, C. Computational methods for transcriptome annotation and quantification using RNA-seq. Nat. Methods 8, 469–77 (2011).

105. Trapnell, C. et al. Transcript assembly and quantification by RNA-Seq reveals unannotated transcripts and isoform switching during cell differentiation. Nat. Biotechnol. 28, 511–515 (2010).

106. Mapleson, D., Venturini, L., Kaithakottil, G. & Swarbreck, D. Efficient and accurate detection of splice junctions from RNA-seq with Portcullis. Gigascience 7, 1–11 (2018).

107. Venturini, L., Caim, S., Kaithakottil, G. G., Mapleson, D. L. & Swarbreck, D. Leveraging multiple transcriptome assembly methods for improved gene structure annotation. Gigascience 7, 1–15 (2018).

108. Fiddes, I. T. et al. Comparative Annotation Toolkit (CAT) - simultaneous clade and personal genome annotation. Genome Res. 231118 (2018) doi:10.1101/231118.

109. Warton, D. I., Duursma, R. A., Falster, D. S. & Taskinen, S. smatr 3- an R package for estimation and inference about allometric lines. Methods Ecol. Evol. (2012) doi:10.1111/j.2041-210X.2011.00153.x.

110. Csürös, M. Malin: Maximum likelihood analysis of intron evolution in eukaryotes. Bioinformatics 24, 1538–1539 (2008).

111. Derelle, R., Philippe, H. & Colbourne, J. K. Broccoli: Combining phylogenetic and network analyses for orthology assignment. Mol. Biol. Evol. 37, 3389–3396 (2020).

112. Eddy, S. R. Accelerated profile HMM searches. PLoS Comput. Biol. 7, (2011).

113. Bernardes, J., Zaverucha, G., Vaquero, C. & Carbone, A. Improvement in Protein Domain Identification Is Reached by Breaking Consensus, with the Agreement of Many Profiles and Domain Co-occurrence. PLoS Comput. Biol. 12, 1–39 (2016).

114. Das, S. et al. CATH FunFHMMer web server: Protein functional annotations using functional family assignments. Nucleic Acids Res. 43, W148–W153 (2015).

115. Dawson, N. L. et al. CATH: An expanded resource to predict protein function through structure and sequence. Nucleic Acids Res. 45, D289–D295 (2017).

116. Sillitoe, I. et al. CATH: Comprehensive structural and functional annotations for genome sequences. Nucleic Acids Res. 43, D376–D381 (2015).

117. Gadagkar, S. R., Rosenberg, M. S. & Kumar, S. Inferring species phylogenies from multiple genes: Concatenated sequence tree versus consensus gene tree. J. Exp. Zool. Part B Mol. Dev. Evol. 304, 64–74 (2005).

118. Seo, T. K., Kishino, H. & Thorne, J. L. Incorporating gene-specific variation when inferring and evaluating optimal evolutionary tree topologies from multilocus sequence data. Proc. Natl. Acad. Sci. U. S. A. 102, 4436–4441 (2005).

119. Zhang, C., Rabiee, M., Sayyari, E. & Mirarab, S. ASTRAL-III: Polynomial time species tree reconstruction from partially resolved gene trees. BMC Bioinformatics 19, 15–30 (2018).

120. Pease, J. B., Brown, J. W., Walker, J. F., Hinchliff, C. E. & Smith, S. A. Quartet Sampling distinguishes lack of support from conflicting support in the green plant tree of life. Am. J. Bot. 105, 385–403 (2018).

121. Yang, Z. PAML 4: Phylogenetic analysis by maximum likelihood. Mol. Biol. Evol. (2007) doi:10.1093/molbev/msm088.

122. Kumar, S., Stecher, G., Suleski, M. & Hedges, S. B. TimeTree: A Resource for Timelines, Timetrees, and Divergence Times. Mol. Biol. Evol. 34, 1812–1819 (2017).

123. Rambaut, A. & Drummond, A. J. Tracer v14, Available from http://beast.bio.ed.ac.uk/Tracer. (2007).

124. Malinsky, M., Matschiner, M. & Svardal, H. Dsuite - Fast D-statistics and related admixture evidence from VCF files. Mol. Ecol. Resour. 21, 584–595 (2021).

125. Larkin, M. a. et al. Clustal W and Clustal X version 2.0. Bioinformatics 23, 2947–2948 (2007).

126. Curran, D. M., Gilleard, J. S. & Wasmuth, J. D. MIPhy: Identify and quantify rapidly evolving members of large gene fam. PeerJ 2018, 1–17 (2018).

127. Wertheim, J. O., Murrell, B., Smith, M. D., Kosakovsky Pond, S. L. & Scheffler, K. RELAX: Detecting relaxed selection in a phylogenetic framework. Mol. Biol. Evol. 32, 1–13 (2014).

128. Cicconardi, F., Marcatili, P., Arthofer, W., Schlick-Steiner, B. C. & Steiner, F. M. Positive diversifying selection is a pervasive adaptive force throughout the Drosophila radiation. Mol. Phylogenet. Evol. 112, 230–243 (2017).

129. Cicconardi, F. et al. Chemosensory adaptations of the mountain fly Drosophila nigrosparsa (Insecta: Diptera) through genomics’ and structural biology’s lenses. Sci. Rep. 7, 43770 (2017).

130. Whelan, S., Irisarri, I. & Burki, F. PREQUAL: Detecting nonhomologous characters in sets of unaligned homologous sequences. Bioinformatics 34, 3929–3930 (2018).

131. Di Franco, A., Poujol, R., Baurain, D. & Philippe, H. Evaluating the usefulness of alignment filtering methods to reduce the impact of errors on evolutionary inferences. BMC Evol. Biol. 19, 1–17 (2019).

132. Kosakovsky Pond, S. L. et al. A random effects branch-site model for detecting episodic diversifying selection. Mol. Biol. Evol. 28, 3033–3043 (2011).

133. Smith, M. D. et al. Less is more: an adaptive branch-site random effects model for efficient detection of episodic diversifying selection. Mol. Biol. Evol. 32, 1342–1353 (2015).

134. Kosakovsky Pond, S. L. et al. HyPhy 2.5 - A Customizable Platform for Evolutionary Hypothesis Testing Using Phylogenies. Mol. Biol. Evol. 37, 295–299 (2020).

135. Falcon, S. & Gentleman, R. Using GOstats to test gene lists for GO term association. Bioinformatics 23, 257–258 (2007).

136. Klopfenstein, D. V. et al. GOATOOLS: A Python library for Gene Ontology analyses. Sci. Rep. 8, 1–17 (2018).

137. Zuberi, K. et al. GeneMANIA prediction server 2013 update. Nucleic Acids Res. 41, 115–122 (2013).

138. Montojo, J., Zuberi, K., Rodriguez, H., Bader, G. D. & Morris, Q. GeneMANIA: Fast gene network construction and function prediction for Cytoscape. F1000Research 3, 1–8 (2014).

139. Vlasblom, J. et al. Novel function discovery with GeneMANIA: A new integrated resource for gene function prediction in Escherichia coli. Bioinformatics 31, 306–310 (2014).

140. Siepel, A. et al. Evolutionarily conserved elements in vertebrate, insect, worm, and yeast genomes. Genome Res. 15, 1034–1050 (2005).

141. Hubisz, M. J., Pollard, K. S. & Siepel, A. PHAST and RPHAST: phylogenetic analysis with space/time models. Brief. Bioinform. 12, 41–51 (2011).

142. Emanuelsson, O., Brunak, S., von Heijne, G. & Nielsen, H. Locating proteins in the cell using TargetP, SignalP and related tools. Nat. Protoc. 2, 953–71 (2007).

143. Söding, J., Biegert, A. & Lupas, A. N. The HHpred interactive server for protein homology detection and structure prediction. Nucleic Acids Res. 33, 244–248 (2005).

144. Baek, M. et al. Accurate prediction of protein structures and interactions using a three-track neural network. Science (80-.). 373, 871–876 (2021).

145. Ruiz-Serra, V. et al. Assessing the accuracy of contact and distance predictions in CASP14. Proteins Struct. Funct. Bioinforma. 89, 1888–1900 (2021).

146. Jr., F. J. M. The Kolmogorov-Smirnov Test for Goodness of Fit. J. Am. Stat. Assoc. 46, 68–78 (1951).

